# Combined BRAF and RAS inhibition overcomes acquired resistance in *BRAF*-mutated colorectal cancer

**DOI:** 10.64898/2026.11.24.768533

**Authors:** Kushtrim Kryeziu, Jakob M. Stenersen, Luís Nunes, Max Z. Totland, Ida Sophie Gjøstøl Strømsvåg, Nora S. Strømnes, Vilde C. W. Elster, Barbara Niederdorfer, Visame Jonuzi, Edward Leithe, Ina A. Eilertsen, Ragnhild A. Lothe, Anita Sveen

**Affiliations:** Department of Molecular Oncology, Institute for Cancer Research, Oslo University Hospital, Oslo, Norway; Institute for Clinical Medicine, Faculty of Medicine, University of Oslo, Oslo, Norway; Department of Biosciences, Faculty of Mathematics and Natural Sciences, University of Oslo, Oslo, Norway

**Author notes:** Corresponding authors: Anita Sveen and Kushtrim Kryeziu,; Address: Department of Molecular Oncology, Institute for Cancer Research Oslo University Hospital HE - Norwegian Radium Hospital, Mail address: P. O. Box 4953 Nydalen, NO-0424 Oslo, Norway.

**Keywords:** CRC, BRAF, RAS, drug resistance, cell line, patient-derived organoid, pharmacogenomics

## Abstract

*BRAF*^V600E^-mutated colorectal cancer is aggressive and associated with poor patient prognosis. Combined BRAF and EGFR inhibition improves patient outcomes but acquired resistance inevitably develops and limits treatment efficacy. We integrated molecular profiling, long-term drug exposure and drug combination screening of *BRAF*-mutated cell lines (n = 17) and patient-derived organoids (n = 17) to identify and target resistance mechanisms. Intrinsic and acquired resistance to dual BRAF and EGFR inhibition converged on several shared transcriptomic features, including IGF1R, PI3K, inflammatory, and proteasome-associated signaling. Screening of 28 drug combinations revealed context-dependent sensitivity to co-inhibition of BRAF with IGF1R, MEK and SRC inhibitors. Acquired resistance was additionally associated with strong RAS activation, including emerging *NRAS* hotspot mutations, *KRAS* amplification and transcriptomic MAPK signaling. RAS inhibitors daraxonrasib and RMC-7977 showed pronounced synergy with encorafenib in models with acquired RAS mutations, restoring treatment sensitivity and suppressing ERK1/2 phosphorylation. These findings identify acquired resistance associated with RAS reactivation as a therapeutic vulnerability and provide a preclinical rationale for combined BRAF and RAS inhibition in *BRAF*^V600E^-mutated colorectal cancer.

## Introduction

Colorectal cancer (CRC) remains a leading cause of cancer-related mortality worldwide despite advances in systemic therapy and molecular stratification (1). The *BRAF*^V600E^ mutation is found in approximately 8-12% of CRCs and identifies a distinct and particularly aggressive subgroup characterized by rapid disease progression, limited response to chemotherapy, and poor patient survival (2,3).

The *BRAF*^V600E^ hotspot mutation drives constitutive activation of the mitogen-activated protein kinase (MAPK) signaling pathway and functions as a dominant oncogenic driver across multiple cancer types. Although *BRAF*^V600E^ is a potent drug target, BRAF inhibitor monotherapy is ineffective in CRC due to rapid adaptive feedback reactivation of MAPK signaling through EGFR and RAS proteins, restoring downstream phosphorylation of ERKs (4). These mechanistic insights have informed combination treatment strategies for *BRAF*^V600E^-mutated metastatic CRC. The randomized phase III trial BEACON CRC promoted dual BRAF and EGFR inhibition with encorafenib plus cetuximab as a standard of care in the treatment refractory setting, demonstrating improved overall survival and response rates compared with chemotherapy plus cetuximab (5). More recently, the randomized phase III trial BREAKWATER extended this targeted combination strategy to the first-line setting, demonstrating improved overall and progression-free survival with the addition of encorafenib plus cetuximab to standard chemotherapy with mFOLFOX6 or FOLFIRI (2,6,7). Together, these trials illustrate the translational impact of mechanistically informed treatment design.

Despite these advances, resistance to BRAF-targeted combination therapies is a major clinical challenge in CRC. Intrinsic and acquired resistance limit the response rate and duration of response, and virtually all patients ultimately experience disease progression (8–10). Analyses of circulating tumor DNA in post-progression liquid biopsies have identified restoration of MAPK signaling as a dominant mechanism of acquired resistance (8). Resistance-associated alterations include activating hotspot mutations in *KRAS*, *NRAS and MAP2K1*, as well as amplification of *KRAS, BRAF*, and receptor tyrosine kinases such as *MET*. Multiple resistance alterations may co-occur within the same tumor (8). These findings emphasize the roles of tumor heterogeneity and clonal evolution in shaping therapeutic response, and support longitudinal molecular profiling to capture the dynamic landscapes of treatment resistance.

Preclinical studies have provided important mechanistic insights and guided the development of rational combination therapies in *BRAF*^V600E^-mutated CRC. Beyond feedback reactivation of MAPK signaling, activation of parallel pathways such as PI3K-mTOR signaling can also limit therapeutic efficacy, and combined pathway blockade is required for effective treatment (11). Experimental models of acquired resistance to combined BRAF and EGFR inhibition have revealed recurrent restoration of MAPK pathway activity through RAS-RAF-MEK-ERK signaling, supporting vertical pathway inhibition as a strategy to overcome resistance (9). SRC-driven inflammatory signaling has also been identified as a reversible resistance program (12). Together, these studies show that resistance can emerge through both adaptive signaling responses and acquired molecular alterations in a context-dependent manner, highlighting the need for systematic functional testing to prioritize therapeutically actionable vulnerabilities.

In this study, we performed integrated molecular profiling and drug combination screening of *BRAF*^V600E^-mutated CRC cell lines and patient-derived organoids (PDOs) with intrinsic and acquired resistance to BRAF-targeted therapy. We aimed to define resistance mechanisms and identify drug combinations capable of restoring treatment sensitivity in resistant models.

## Results

### Preclinical models and transcriptomic associations of BRAF inhibitor sensitivity

Preclinical models of *BRAF*-mutated CRC were identified from a panel of 103 conventional cell lines (13) and 213 PDOs from 102 patients (15). Of these, 17 (16.5%) cell lines and 17 (8%) PDOs from five patients (5%) had a confirmed *BRAF* class I hotspot (p.V600E or p.V600K) mutation (Fig. 1A-B and Table S1). Most *BRAF* class I mutated cell lines (82.4%) were established from primary tumors and all PDOs from resected liver metastases (n = 2-5 tumors per patient). All *BRAF*^V600^ models were *KRAS*/*NRAS* (RAS) wild-type, and most of the cell lines (64.7%) and all PDOs were microsatellite stable (MSS). Additional frequent alterations in key CRC-associated genes included *APC* and *TP53* mutations, as well as *PIK3CA* mutations in a subset (Fig. 1A-B and Table S1).

**Figure 1.**
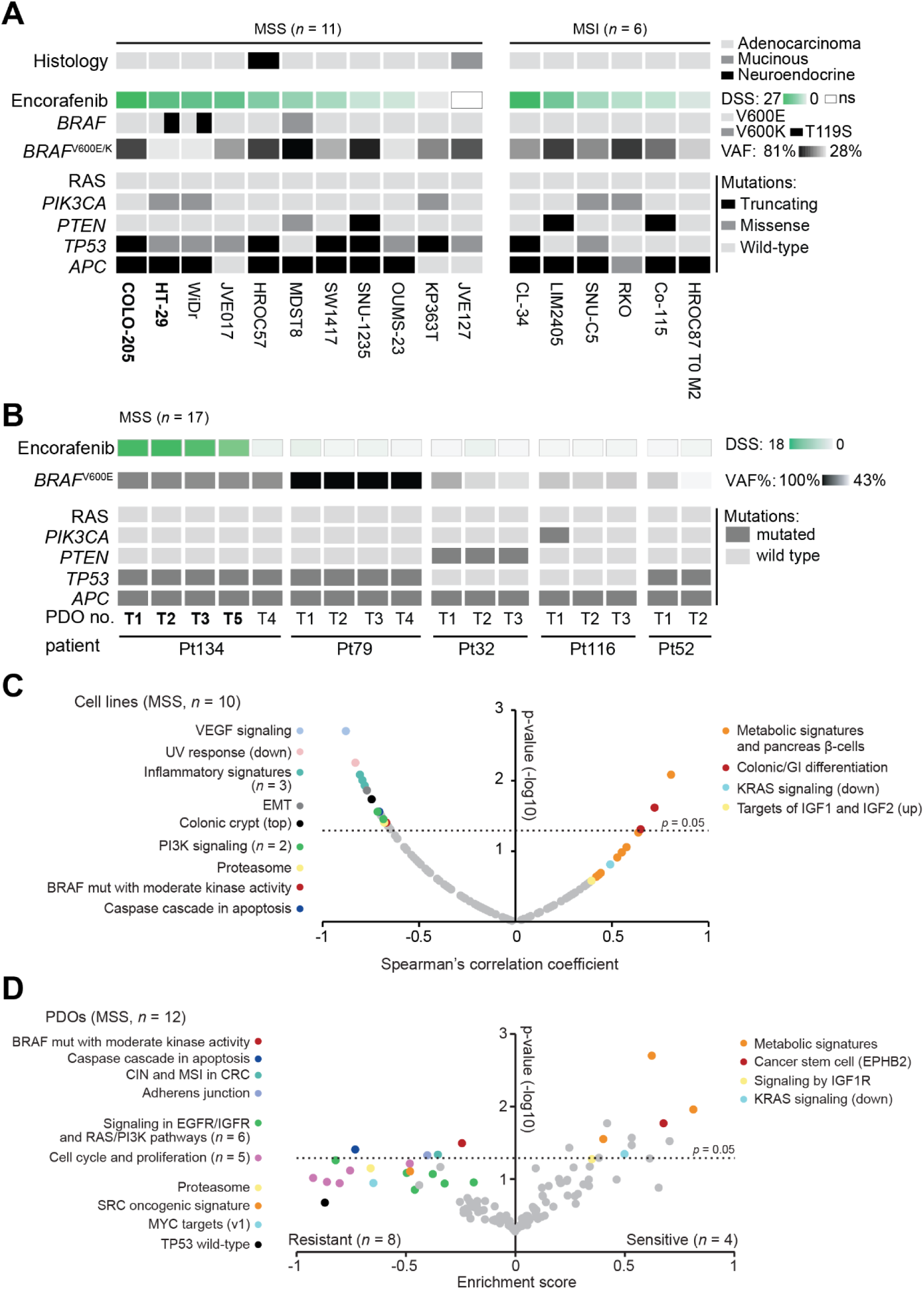
Overview and transcriptomic profiles of the *BRAF*^V600^-mutated cell lines and PDOs. Histological and molecular characteristics of cell lines (**A**) and PDOs (**B**) ordered by MSI status and encorafenib drug sensitivity scores (DSS), illustrating co-occurring mutations in RAS (*KRAS* and *NRAS*), *PIK3CA*, *PTEN*, *TP53* and/or *APC*. Additional annotations indicate cell line histology, variant allele frequency (VAF) of *BRAF*^V600E/K^, and PDOs derived from multiple metastatic lesions of each of 5 patients. Volcano plots of (**C**) Spearman correlations between single-sample enrichment scores of a custom gene set collection (*n* = 124 gene sets) and encorafenib sensitivity in MSS *BRAF*^V600E/K^-mutated cell lines and (**D**) enrichment scores from gene set enrichment analyses comparing encorafenib-sensitive (DSS > 5) and encorafenib-resistant PDOs (DSS ≤ 5). Dashed lines indicate the threshold for statistical significance, and selected gene sets are highlighted with color keys.

Four BRAF inhibitors, agerafenib, dabrafenib, encorafenib, and vemurafenib, were evaluated for anti-cancer activity across all cell lines as part of a high-throughput drug screen (14). This confirmed specific sensitivity among *BRAF*^V600E/K^-mutated models, irrespective of microsatellite instability status (Fig. S1A-B; JVE127 was not scored). The activities of the BRAF inhibitors were strongly correlated (Pearson correlation coefficients > 0.83; Fig. S1D), and encorafenib was selected for subsequent studies based on its high concordance with the other BRAF inhibitors and its established clinical use in *BRAF*^V600E^-mutated CRC. The proportion of sensitive models (drug sensitivity score, DSS > 5) was lower among PDOs than cell lines (4 of 17 PDOs, 23% versus 12 of 16 cell lines, 75%; p = 0.005 from Fisher’s exact test; Fig. S1B-C), suggesting a different drug response pattern to single-agent encorafenib between monolayer cell line cultures and three-dimensional PDOs. Drug response was not associated with the variant allele frequency of *BRAF*^V600^ (Fig. 1A-B).

Among the MSS *BRAF*^V600^-mutated cell lines (n = 10), encorafenib sensitivity was correlated to transcriptomic single-sample enrichment scores of several gene sets in a custom collection (n = 124 gene sets; Table S2). Sensitive cell lines showed enrichment with metabolic signatures, whereas resistant cell lines showed enrichment with signatures related to inflammation, PI3K signaling, and proteasome activity (Fig. 1C and Table S2). Gene set enrichment analysis comparing sensitive and resistant PDOs (n = 12 samples with available gene expression data; Fig. S2A) showed similar trends, including enrichment with EGFR-RAS, IGF, and PI3K-related signatures in resistant models, although these did not reach statistical significance (Fig. 1D and Table S2). In the patient represented by both sensitive and resistant PDOs (n = 4 and n = 1, respectively), a within-patient comparison showed similar enrichment patterns (Fig. S2B).

None of the compounds evaluated across the complete pharmacological screens (n = 620 and 51 drugs in cell lines and PDOs, respectively) showed greater selective activity against *BRAF*^V600E^-mutated models than the BRAF inhibitors (Fig. S2C-D). Some sensitivity to PI3K-mTOR pathway inhibition was observed in both model systems, including to gedatolisib in cell lines (Wilcoxon rank-sum test, adjusted p = 0.02) and everolimus in PDOs (adjusted p = 0.02; Table S3). Mutated models were largely resistant to inhibitors targeting upstream components of the EGFR-MAPK pathway or parallel pathways, including EGFR/HER2 (afatinib and lapatinib) and SRC family kinases (dasatinib). Mutated PDOs were also resistant to the IGF1R inhibitor BMS-754807 and the MEK inhibitor binimetinib (Table S3), consistent with the transcriptomic profiles of intrinsic resistance. Triple inhibition of the EGFR-MAPK pathway with afatinib, encorafenib and trametinib (MEK1/2 inhibitor) was tested in PDOs at equimolar concentrations (1:1:1 ratio) and showed significantly higher DSS than each single agent alone (Fig. S2E). As previously shown (15), this effect was observed in both *BRAF*^V600E^ and wild-type PDOs (paired Wilcoxon tests, all p < 0.01), although the increase in sensitivity relative to encorafenib alone was higher in the wild-type group (Wilcoxon rank-sum test, p = 0.005).

### Generation of cell line models with acquired resistance

To investigate the mechanisms of acquired resistance to BRAF inhibition in CRC, we established resistance models to encorafenib alone and in combination with afatinib using the two most sensitive MSS cell lines, HT-29 and COLO-205 (Fig. 1A). Parental cells were continuously exposed to progressively increasing drug concentrations until stable proliferation was maintained under treatment (Fig. 2A-B; Fig. S3). Resistance was defined as sustained viability at ≥ 1 µM encorafenib and ≥ 1 µM afatinib, which far exceeds the half-maximal inhibitory concentration (IC₅₀) of the parental lines (encorafenib IC₅₀: 6.6 nM and 3.31 nM in HT-29 and COLO-205, respectively; afatinib as single agent was not effective in any of the models; Fig. 2C). Stable resistance to encorafenib emerged after 140 and 160 days in HT-29^E^ and COLO-205^E^ cells, respectively, with IC₅₀ values exceeding the highest concentration tested (1 µM) after 72 h of drug incubation (Fig. S3A, D). Two approaches were used to generate resistance to combined encorafenib plus afatinib: (i) simultaneous selection by gradual dose escalation of both drugs in parental cells (generating HT-29^AE^ and COLO-205^AE^ sublines; Fig. S3B, E); and (ii) sequential selection with further exposure of encorafenib-resistant sublines (HT-29^E^ and COLO-205^E^) to increasing concentrations of the drug combination (generating HT-29^EAE^ and COLO-205^EAE^ sublines; Fig. S3C, F). Resistance developed after 50 days and 60 days for HT-29^AE^ and COLO-205^AE^ cells, respectively, using the simultaneous-selection approach, and after 56 and 32 days in HT-29^EAE^ and COLO-205^EAE^ cells, respectively, using the sequential-selection approach. No major phenotypic changes were observed in the resistant sublines compared with their parental counterparts (Fig. 2A-B), and the variant allele frequencies of *BRAF*^V600E^ and other baseline mutations remained stable (Fig. S3).

**Figure 2.**
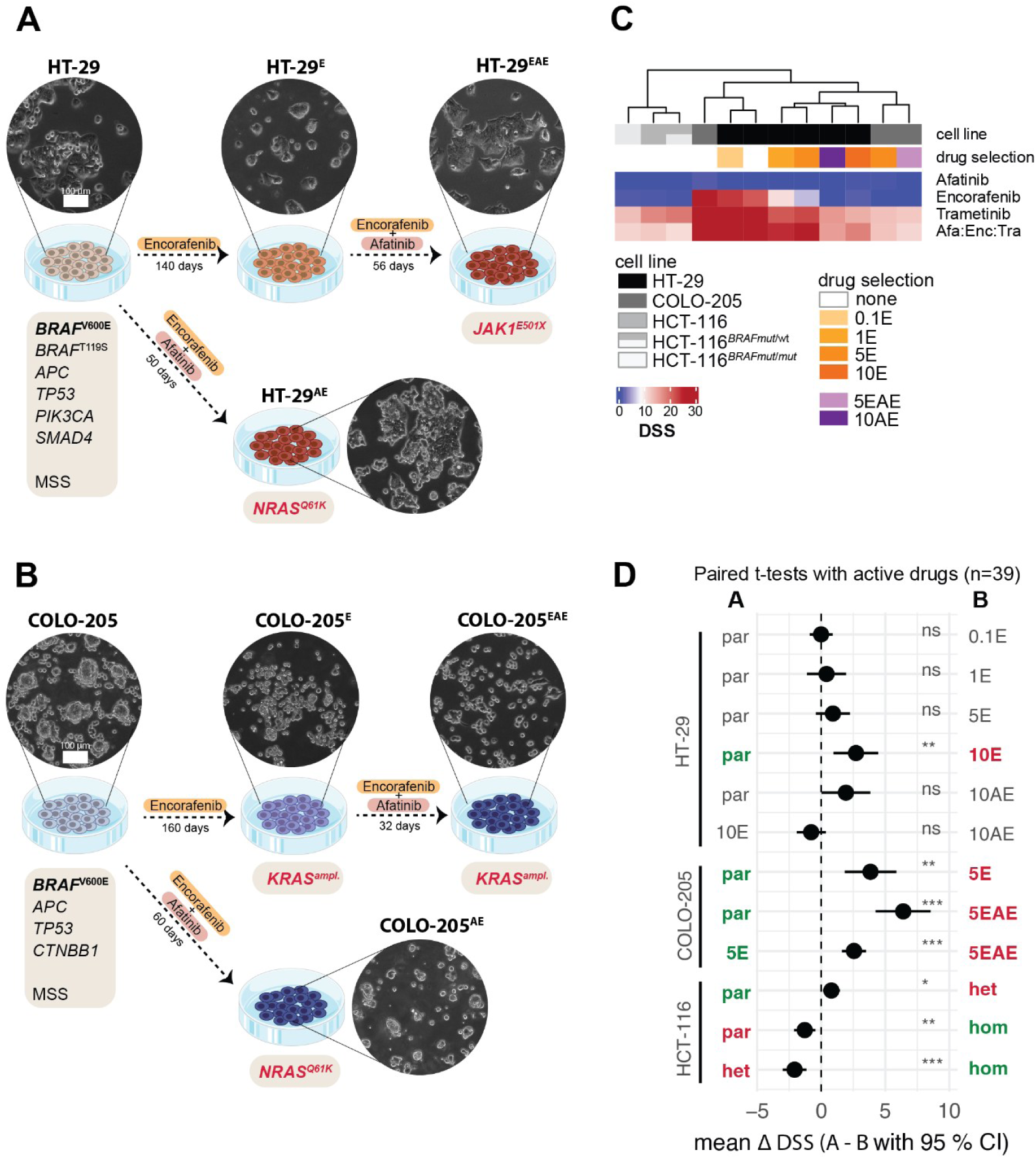
CRC cell line models with acquired resistance to BRAF and EGFR inhibition. Schematic overview of the stepwise generation of (**A**) HT-29 and (**B**) COLO-205 sublines with acquired resistance to encorafenib (HT-29^E^ and COLO-205^E^) or to the combination of encorafenib and afatinib (HT-29^EAE^ and HT-29^EA^; COLO-205^EAE^ and COLO-205^EA^), achieved through drug exposure at incrementally increasing concentrations over the indicated time periods. Representative phase-contrast images are shown for each subline (scale bar, 100 μm). Acquired genetic resistance mutations are indicated in red. **C**) Hierarchical clustering and heatmap of cell lines based on drug sensitivity scores (DSS) across 47 drugs, using Euclidean distance and complete-linkage clustering. Heatmap annotations indicate cell line identity and drug selection conditions. For clarity, the heatmap displays DSS values only for afatinib, encorafenib, trametinib, and the triple combination of afatinib, encorafenib and trametinib (Afa:Enc:Tra; 1:1:1 concentration ratio). **D**) Forest plot showing differences in DSS (ΔDSS) from paired comparisons (A vs B samples) of parental (par) and drug-selected or CRISPR-edited sublines (mean ± 95% confidence intervals; n = 39 drugs). Heterozygous (het) and homozygous (hom) refer to the *BRAF*^V600E^ mutations. Statistical significance was determined by paired t-test; ns, not significant; *p < 0.05; **p < 0.01; ***p < 0.001. Colors indicate significant pairwise comparisons, with green marking the more sensitive model and red marking the more resistant model within each pair.

A drug sensitivity screen (n = 47; Table S4) confirmed acquired resistance to encorafenib and sustained resistance to afatinib in all fully resistant models (HT-29^10E^, HT-29^10AE^, COLO-205^5E^, and COLO-205^5EAE^; Fig. 2C; Fig. S4A). The MEK inhibitor trametinib and the triple combination of encorafenib, afatinib, and trametinib at equimolar concentrations retained activity in HT-29 sublines with resistance acquired at lower encorafenib concentrations (HT-29^0.1E^, HT-29^1E^, HT-29^5E^). However, fully resistant models of both cell lines showed pronounced resistance to the triple MAPK pathway-targeting combination, with a decrease in DSS values of at least 10 relative to the corresponding parental cell lines. There were no acquired vulnerabilities to any of the drugs in the panel (Fig. S4A), and the resistant sublines showed reduced overall drug sensitivity relative to the parental cell lines, quantified as the mean DSS across drugs (39 drugs with DSS > 5; Fig. 2D). The COLO-205 subline with dual afatinib and encorafenib resistance (COLO-205^EAE^) also showed reduced overall drug sensitivity relative to the subline with resistance to encorafenib alone (COLO-205^E^). A corresponding trend was not observed in the HT-29 background, but principal component analysis of drug sensitivities showed that samples clustered according to acquired resistance phenotypes rather than cell line identity, supporting the emergence of a convergent multi-drug resistance profile in both cell line backgrounds (Fig. S4B).

### Genetic and transcriptomic mechanisms of acquired resistance

Mutation profiling of 20 CRC-relevant genes identified acquired RAS mutations across four of the six resistant HT-29 and COLO-205 sublines (Fig. 2A-B). *NRAS*^Q61K^ (c.181C>A) mutations emerged in both sublines selected under combined BRAF and EGFR inhibition (HT-29^AE^ and COLO-205^AE^). Additionally, *KRAS* amplification was detected in COLO-205^E^ following encorafenib selection and increased further after subsequent selection with encorafenib and afatinib, with estimated DNA copy numbers of 23 and 46 in COLO-205^E^ and COLO-205^EAE^, respectively (Fig. S3D, F; Fig. S5A). The amplification was associated with a substantial increase in *KRAS* expression (Fig. S5B). The HT-29^E^ subline acquired a truncating *JAK1*^E501X^ (c.1501G>T) mutation at an intermediate stage of encorafenib resistance. This mutation was further selected during combined encorafenib and afatinib exposure, with increasing variant allele frequency from 6% in HT-29^E^ to 30% in HT-29^EAE^ (Fig. S3A, C).

The involvement of RAS alterations in encorafenib resistance was further supported by drug screening of the *KRAS*^G13D^-mutated HCT-116 cell line and isogenic CRISPR-engineered sublines harboring either a heterozygous or homozygous *BRAF*^V600E^ mutation (obtained from GeneCopoeia, Inc.; Fig. S4C). Consistent with the *KRAS*-mutated background, parental cells were resistant to afatinib and the engineered *BRAF* mutation did not confer sensitivity to encorafenib or the afatinib-encorafenib-trametinib triple combination (Fig. 2D and Fig. S4A).

Transcriptomic profiling and single-sample enrichment analysis of the custom gene set collection in HT-29 and COLO-205 sublines (Table S2) showed that acquired resistance to both single-agent encorafenib and the combination with afatinib was associated with increased enrichment scores of EGFR-RAS-PI3K and IGF-IGF1R-RAS-ERK pathway activity, consistent with the acquired RAS mutations (Fig. 3A, C). Signatures of oxidative phosphorylation and the proteasome were also enriched after dual resistance development (Fig. 3D). In contrast, signatures of IGF1/IGF2 targets, PI3K activity, and unfolded protein response were depleted in resistant models. Several signatures of the SRC and IGF pathways were involved in resistance development, including both enrichment and depletion in relation to the parental cell lines. Analyses of HCT-116 sublines with engineered *BRAF* mutations indicated that similar pathways were also involved in intrinsic resistance (Fig. S4D).

**Figure 3.**
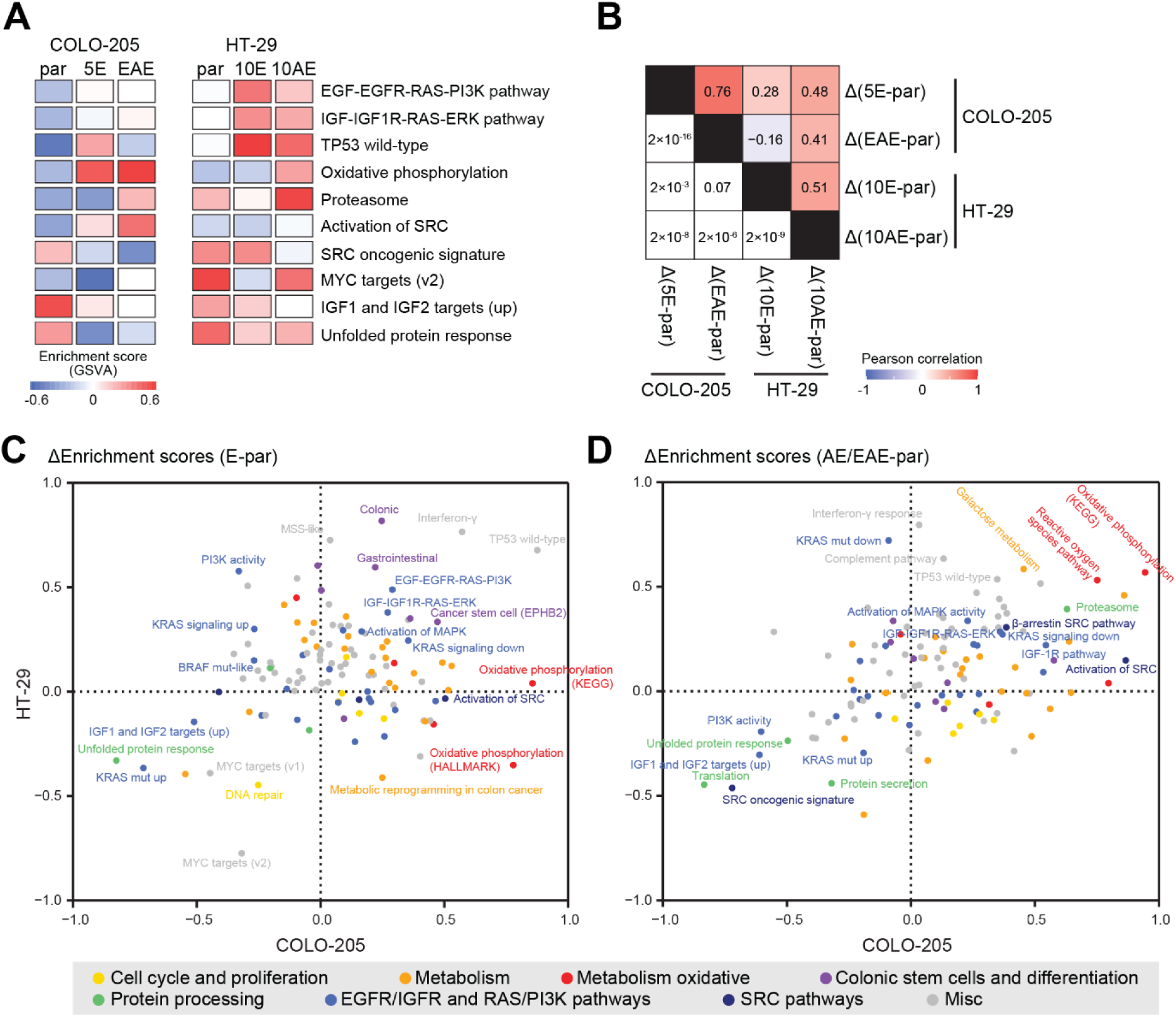
Transcriptomic profiles associated with acquired resistance to BRAF inhibition and BRAF-EGFR combination therapy. **A**) Single-sample enrichment scores of selected gene sets (n = 10) showing the largest differences between parental (par) COLO-205 and HT-29 cells and their derived sublines with acquired resistance to encorafenib (E) or the combination of encorafenib and afatinib (EAE/EA). **B**) Correlation matrix of Pearson correlation coefficients (upper triangle) and corresponding *p*-values (lower triangle) representing differences (Δ) in single-sample enrichment scores across the custom gene set collection (n = 124 gene sets) between parental cells and models with acquired resistance. Scatter plots of differences in single-sample enrichment scores (n = 124 gene sets) between parental cells and (**C**) models with acquired resistance to single-agent encorafenib or (**D**) models with acquired resistance to the combination of encorafenib and afatinib.

Overall, transcriptomic changes associated with acquired resistance to the combination treatment were largely consistent between HT-29 and COLO-205, as indicated by correlation of the differences in gene set enrichment scores between resistant and parental cells (Pearson r = 0.41, *p* = 2×10^-6^), although some cell line-specific alterations were observed (Fig. 3B, D). In contrast, transcriptomic changes associated with resistance to single-agent encorafenib were more heterogeneous between the two cell lines (Pearson r = 0.28, *p* = 0.002).

### Drug combinations targeting transcriptomic resistance mechanisms

A drug combination screen was designed using a BRAF inhibitor, primarily encorafenib, as the anchor drug paired with 22 additional drugs in full 7×7 concentration matrices (Table S5). The screen was used to functionally evaluate candidate mechanisms of acquired resistance identified in the HT-29 and COLO-205 sublines and intrinsic resistance inferred from parental cell lines and CRISPR engineered HCT-116 models. The combination screen was evaluated in nine cell lines, including two encorafenib sensitive (parental HT-29 and COLO-205) and seven resistant (HT-29 and COLO-205 sublines and the engineered HCT-116 models), as well as in four PDOs, including two sensitive and two with intrinsic resistance to encorafenib (Fig. 4A). Combination treatments were generally less effective in resistant than sensitive models, as reflected by lower mean activity (percent decrease in viability; Wilcoxon test, p = 2.19 × 10⁻²⁶) and weaker synergy (mean Bliss scores; Wilcoxon rank-sum test, p = 7 × 10⁻⁴; Fig. S6A-C). However, models with resistance to single-agent encorafenib (HT-29^E^ and COLO-205^E^) showed stronger synergistic response to combinations of BRAF and EGFR inhibitors than the corresponding parental cell lines (Fig. S6A-B; Fig. S7). This combination was also highly synergistic in all PDOs, but models with dual drug resistance showed antagonistic response between BRAF and EGFR inhibitors, consistent with the acquired *NRAS*^Q61K^ mutation in HT-29^AE^ and *KRAS* amplification in COLO-205^EAE^, and the intrinsic *KRAS*^G13D^ mutation in HCT-116 sublines (Fig. 4A; Fig. S7). In contrast, the dual resistance models showed synergistic response to co-inhibition of BRAF and its downstream mediator MEK1/2 (encorafenib-trametinib), consistent with elevated transcriptomic pathway activity and enhanced RAS-BRAF-MEK signaling (Fig. 4A; Fig. S8) (28). Synergy between encorafenib and the PI3K/mTOR inhibitor gedatolisib was observed in parental HT-29 and COLO-205 cells and in PDOs, consistent with the transcriptomic features associated with intrinsic resistance (Fig. 1C, D; Fig. S9) and with established mechanisms of resistance to BRAF inhibition (8,11).

**Figure 4.**
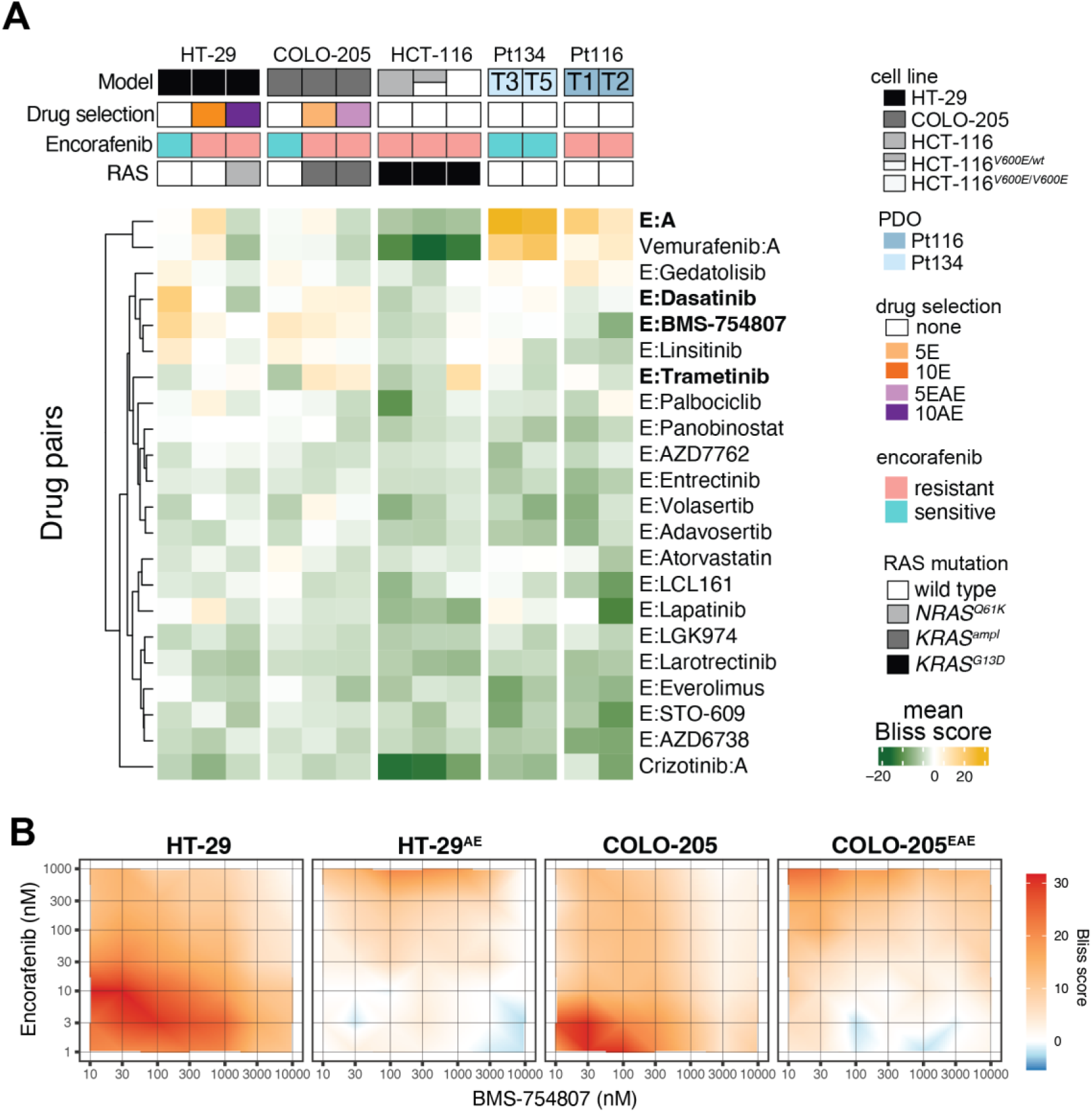
Synergy screening of encorafenib and drugs targeting pathways of transcriptomic resistance mechanisms. **A**) Heatmap of mean Bliss synergy scores for encorafenib (E) combined with 20 inhibitors, vemurafenib plus afatinib (A) and crizotinib plus afatinib (A) across CRC cell lines (n = 9) and PDO models (n = 4). Columns represent samples and rows represent drug pairs ordered by hierarchical clustering of the mean Bliss scores using complete linkage. Annotation bars indicate cell lines, resistance status to encorafenib, drug selection background, and RAS mutation status. **B**) Bliss synergy matrices of the full 7x7 drug combination grid of encorafenib and the IGF1R inhibitor BMS 754807 in parental HT-29 and COLO-205 cell lines and corresponding sublines with dual encorafenib-afatinib resistance. Positive Bliss scores indicate synergy, whereas negative scores indicate antagonism.

Beyond these well-known synergistic activities, the combination of encorafenib with IGF1R inhibitors (particularly BMS-754807) exhibited consistent synergistic responses across cell lines, including in models with acquired resistance and in the HCT-116 subline with homozygous *BRAF*^V600E^ mutation (Fig. 4A; Fig. S6A-B; Fig. S10-S11). However, synergy was most pronounced in parental HT-29 and COLO-205 cells at lower encorafenib concentrations (Fig. 4B; Fig. S10-S11). These findings were consistent with transcriptomic evidence implicating IGF pathway activity in both acquired and intrinsic resistance to BRAF inhibition.

The combination of encorafenib with dasatinib, an inhibitor of ABL, SRC and c-KIT kinases, also showed synergistic activity in parental HT-29 and COLO-205 cells, but distinct and opposing response patterns across the corresponding resistant sublines (Fig. 4A). Strongest synergistic response was observed in parental HT-29 cells (Fig. S12) and acquired resistance sublines of COLO-205. For improved target specificity, the selective SRC inhibitor eCF506 was combined with encorafenib as part of a second combination screen in HT-29 and HT-29^AE^ cells (Table S6). This supported greater activity in the parental than the resistant subline, with mean viability inhibition of 51% and 17%, respectively, and mean Bliss scores of 1.26 and -5.3, indicating additive and antagonistic interactions, respectively (Fig. S13B, E). The response pattern to the encorafenib-dasatinib combination in COLO-205 was consistent with increased transcriptomic SRC activity in resistant sublines, but the combined findings for SRC inhibition indicated that this resistance mechanism is context dependent (Fig. 3A; Fig. S12).

Combinations of encorafenib with the two proteasome inhibitors bortezomib and carfilzomib in HT-29 and HT-29^AE^ cells showed additive or antagonistic effects even at the lowest concentrations of the proteasome inhibitors. These results were not consistent with the strong transcriptomic enrichment of proteasome-related signatures in the acquired resistance models (Fig. 3A, D) and likely reflected the high single-agent activity of the proteasome inhibitors in both parental and resistant HT-29 cells (Fig. S13A, D).

### Efficacy of RAS inhibition in models with acquired resistance

Given the frequent emergence of acquired RAS mutations after combined BRAF and EGFR inhibition, both in patients with CRC (8) and in our preclinical models, we evaluated encorafenib in combination with two panRAS(ON) inhibitors and a SHOC2–RAS protein-protein interaction inhibitor targeting upstream RAS signaling (Table S6). The SHOC2-RAS interaction inhibitor showed low or undetectable single-agent activity both in parental HT-29 and resistant HT-29^AE^ cells and did not enhance the activity of encorafenib (Fig. S13C, F).

In contrast, the combinations of encorafenib with the panRAS inhibitors daraxonrasib and RMC-7977 demonstrated clear synergistic effects in both cell lines. Parental HT-29 cells showed mean Bliss scores of 15 and 9 for daraxonrasib (Fig. 5A-B) and RMC-7977 (Fig. S14A-B), respectively, while parental COLO-205 cells showed mean Bliss scores of 6 and 9, respectively (Fig. 5C-D; Fig. S14C-D). Nonetheless, the synergy was stronger in the dual resistant HT-29^AE^ and COLO-205^EAE^ sublines across a broader concentration range of both daraxonrasib (mean Bliss score of 54 and 22, respectively) and RMC-7977 (mean Bliss score of 45 and 21, respectively; Fig. 5A, C; Fig. S14A, C). Accordingly, synergistic activity was observed across two distinct genetic mechanisms of acquired resistance, although restoration of sensitivity was more pronounced in the *NRAS*^Q61K^-mutated HT-29^AE^ model than in the *KRAS*-amplified COLO-205^EAE^ model. The maximal viability inhibition in HT-29^AE^ cells was similar to parental HT-29 cells (84% vs 85% for encorafenib–daraxonrasib and 89% vs 85% for encorafenib–RMC-7977, respectively), while COLO-205^EAE^ showed only partial restoration of viability inhibition relative to parental COLO-205 (58% vs 82% and 64% vs 86%, respectively; Fig. 5B, D; Fig. S14B, D).

**Figure 5.**
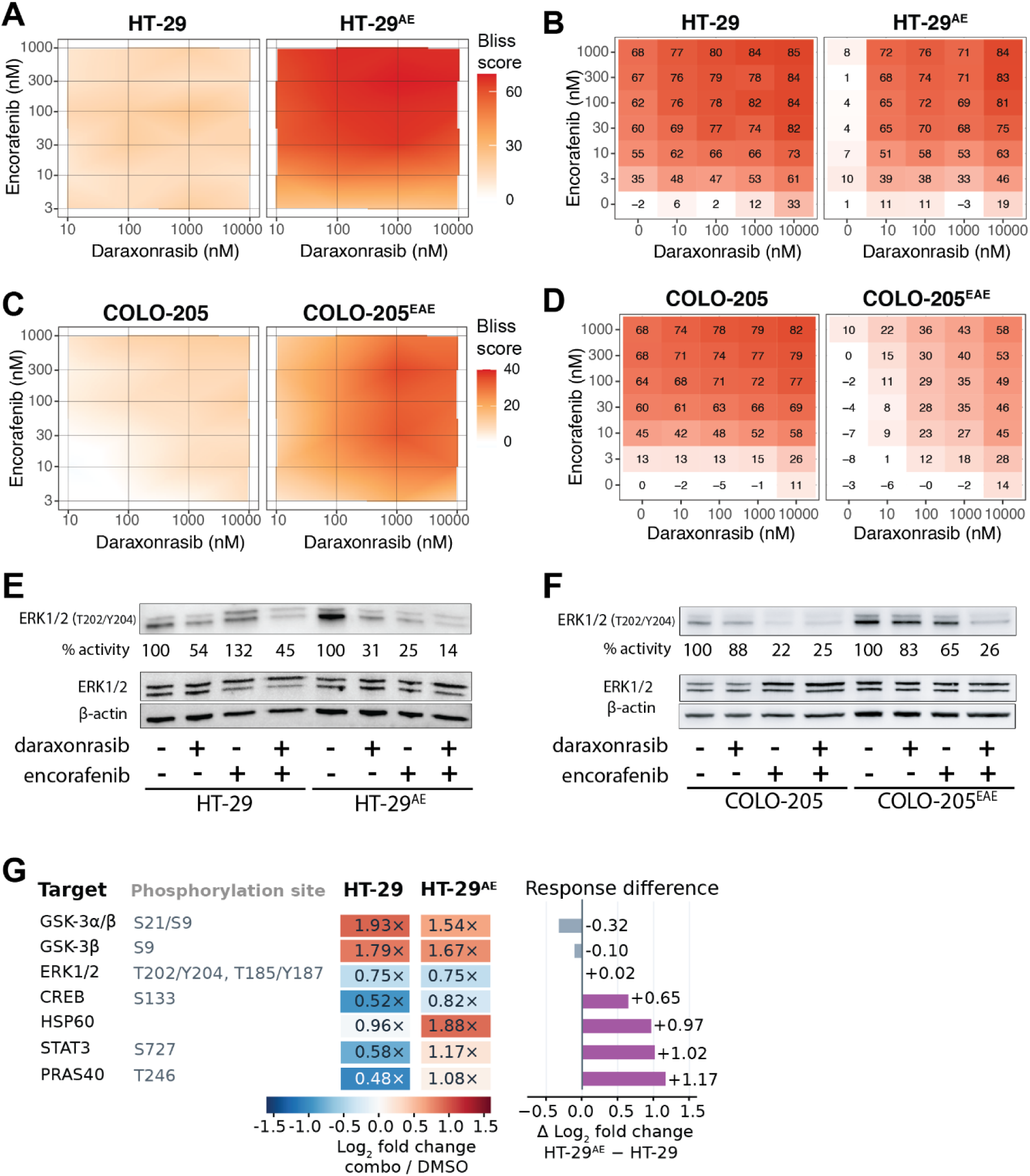
RAS inhibition restores sensitivity to encorafenib in two cell lines with different genetic resistance mechanisms. Bliss synergy matrices (**A** and **C**) and dose-response heatmaps of percent viability inhibition (**B** and **D**) for the combination of encorafenib with the panRAS(ON) inhibitor daraxonrasib in parental HT-29 and COLO-205 cells and their encorafenib and afatinib resistant sublines (HT-29^AE^ and COLO-205^EAE^). (**E-F**) Immunoblot of MAPK pathway activity in the parental HT-29 and COLO-205 cells and resistant HT-29^AE^ and COLO-205^EAE^ sublines after 72h of treatment with DMSO (vehicle control), 1µM encorafenib (BRAF inhibitor), 1µM daraxonrasib (panRAS (ON) inhibitor), or the combination (1µM of each), as indicated. Phosphorylation of ERK1/2 (p-ERK1/2, Thr202/Tyr204) was assessed as a readout of MAPK pathway activity, with total ERK1/2 and β-actin serving as loading controls. ERK1/2 and phospho-ERK1/2 were run on separate membranes. Percent activity values indicate p-ERK1/2 normalized to their corresponding β-actin loading controls and total ERK1/2 levels, with the DMSO-treated condition set to 100% for each cell line. (**G**) Selected protein activities to combined encorafenib and daraxonrasib treatment in parental HT-29 cells and the resistant HT-29^AE^ subline, assessed using a phospho-kinase antibody array. Heatmap colors indicate log₂ fold changes relative to DMSO-treated controls, with the corresponding untransformed fold changes shown within each cell line. Phosphorylation sites are indicated where applicable. Bars show differences in log₂ fold changes between resistant and parental cells (HT-29^AE^ − HT-29), with positive values indicating a greater increase or weaker reduction in the resistant subline.

Consistent with these cell viability results, combined BRAF and RAS inhibition suppressed downstream MAPK signaling in both resistant models, with more pronounced pathway suppression in HT-29^AE^ than in COLO-205^EAE^ cells. Both resistant models had higher ERK1/2 phosphorylation at baseline (DMSO treatment) than the parental lines, and the combination of encorafenib and daraxonrasib achieved greater suppression of ERK1/2 phosphorylation than either drug alone, reducing levels to 14% and 26% of the corresponding DMSO controls in HT-29^AE^ and COLO-205^EAE^ cells, respectively. In parental HT-29 and COLO-205 cells, the combination reduced ERK1/2 phosphorylation to 45% and 25% of the corresponding DMSO controls, respectively (Fig. 5E–F). The effect in parental cells largely reflected single-agent activity, and daraxonrasib reduced ERK1/2 phosphorylation levels to 54% in HT-29 cells and encorafenib to 22% in COLO-205 cells.

Broader phosphorylation profiling of parental HT-29 and resistant HT-29^AE^ cells using a phospho-kinase antibody array revealed largely concordant responses to combined encorafenib and daraxonrasib treatment (37 phosphoproteins; Pearson r = 0.77, p = 2 × 10⁻⁸; Fig. S15A and Table S10). Common responses beyond ERK1/2 (ERK1 T202/Y204; ERK2 T185/Y187) included reduced phosphorylation of S6K (T389, T421/S424), CREB (S133) and p53 (S15, S46 and S392), as well as increased phosphorylation of GSK3 at inhibitory regulatory sites (S9 and S21; Fig. 5G; Fig. S15B-C). Despite this overall concordance, several signaling proteins showed distinct responses in the resistant model. Phosphorylation of AKT (S473) and its downstream substrate PRAS40 (T246) was reduced to 0.43- and 0.48-fold of DMSO controls in parental cells, respectively, but was largely maintained in resistant cells (0.91- and 1.09-fold, respectively). Similarly, CREB (S133) phosphorylation was less strongly suppressed in resistant than parental cells (0.82-versus 0.52-fold). STAT3 (S727) phosphorylation showed opposing responses, decreasing to 0.58-fold in parental cells and increasing to 1.17-fold in resistant cells. Moreover, HSP60 protein abundance increased in resistant cells (1.88-fold) but remained largely unchanged in parental cells (0.96-fold).

Notably, strong synergy between BRAF and RAS inhibition was not observed in the PDO models, none of which harbored RAS mutations. An encorafenib-sensitive PDO showed modest synergy with mean Bliss scores of 9 and 11 for daraxonrasib and RMC-7977, respectively (Fig. S14E-H), similar to the level observed for the parental HT-29 and COLO-205 cells. However, a PDO with intrinsic encorafenib resistance showed no response to either combination. These findings suggest that the combination of BRAF and RAS inhibitors is only beneficial after the development of acquired resistance to BRAF targeted combination therapy and is dependent both on baseline sensitivity to encorafenib and on emergence of RAS-driven signaling.

## Discussion

This study provides a preclinical rationale for targeting acquired resistance mechanisms and restoring sensitivity to BRAF-targeted therapy in CRC. Our preclinical models recapitulated MAPK pathway reactivation as a major mechanism of resistance to combined BRAF and EGFR inhibition and provided experimental evidence that RAS inhibition can overcome this resistance (8,9). Notably, the potential of combined BRAF and RAS inhibition to overcome resistance appeared specific to acquired resistance driven by RAS mutations, but was effective with different mutation types, including *NRAS*^Q61K^ and *KRAS* amplification. These findings are encouraging considering the diversity of genetic mechanisms for RAS protein activation observed in both patients and preclinical models after combined BRAF/EGFR inhibition (8,9). We observed recurrent emergence of *NRAS*^Q61K^ mutations in two different cell lines, and acquired *NRAS* mutation is also the second most common alteration in patients at progression (8). However, another study of 11 CRC cell lines identified a broader diversity of acquired resistance mutations (9), suggesting that the genetic mechanisms underlying MAPK pathway reactivation may be model-specific and influenced by experimental factors, such as the specific treatment regimen used and duration of the treatment pressure. *KRAS* amplification was observed in both the present study and the previous preclinical study, but it is less frequently reported as a resistance mechanism in patients (8). In our study, *KRAS* amplification emerged during the development of resistance to single-agent encorafenib and was retained following subsequent exposure to combined BRAF and EGFR inhibition. This suggests that once established, RAS pathway activation remains a stable driver of resistance under intensified treatment pressure on the MAPK pathway, highlighting the evolutionary robustness of RAS-driven escape mechanisms. From these perspectives, RAS inhibition might be a potent rescue treatment in a substantial proportion of patients with acquired resistance to BRAF-targeted therapy.

Our findings suggest a new clinical indication for RAS inhibition in CRC, which is particularly relevant given recent advances in RAS-targeted drug development (29,30). Several RAS(ON) multi-selective inhibitors have demonstrated broad preclinical activity across RAS-driven cancers, and daraxonrasib has significantly improved progression free survival and overall survival compared to chemotherapy in patients with previously treated RAS-mutant pancreatic cancers (31). In CRC, mutation-specific KRAS^G12C^ inhibitors have shown limited activity as monotherapy, in part because inhibition of KRAS^G12C^ induces adaptive feedback signaling through EGFR and other receptor tyrosine kinases, resulting in activation of MAPK signaling (32). Consistent with this mechanism, clinical studies have shown greater activity of KRAS^G12C^ inhibitors when combined with EGFR blockade (33,34). Clinical data for broader RAS(ON) inhibitors in CRC remain limited, and it is not known whether also this strategy will require targeted combination therapies. Nevertheless, our findings support the further investigation of RAS(ON) inhibitors in CRC and identify acquired resistance to combined BRAF and EGFR inhibition as a new potential therapeutic setting for their use. Additional preclinical and clinical studies are required to determine whether RAS inhibition can overcome resistance and improve the durability of response to combined BRAF/EGFR inhibition therapy *in vivo*.

Despite the restoration of treatment sensitivity *in vitro*, phospho-kinase profiling suggested that cells with acquired resistance retain distinct mechanisms of signaling regulation and stress adaptation during combined BRAF and RAS inhibition. These responses may contribute to cell viability and represent candidate mechanisms of treatment escape, although their functional impact remains to be established.

In our study, transcriptomic mechanisms of intrinsic and acquired resistance to BRAF/EGFR inhibition were largely concordant among models and predominantly involved downstream or parallel signaling nodes to the MAPK pathway, including MEK, PI3K–mTOR, and SRC. Therapeutic targeting of these mechanisms with drug combinations showed efficacy consistent with previous evidence that vertical and parallel pathway inhibition can enhance suppression of MAPK signaling in *BRAF*^V600E^ CRC (8,12). Beyond these mechanisms, we identified synergy between BRAF and IGF1R inhibitors, suggesting that the IGF1R receptor might act as a bypass mechanism to restore MAPK signaling in resistant models and potentially be developed as a therapeutic target in *BRAF*-mutated CRC. This is supported by a preclinical study of resistance to combined BRAF and MEK inhibition in melanoma (35). However, none of these alternative pathways showed comparable efficacy to RAS-inhibition in targeting resistance to BRAF inhibition.

A limitation of this study is the preclinical design, and important determinants of *in vivo* treatment response are not captured, such as immune and stromal interactions in the tumor microenvironment. This is particularly relevant because *BRAF*^V600E^-mutated MSS CRCs can display immune activation, and emerging evidence suggest potential benefit from adding immune checkpoint inhibition to combined BRAF/EGFR inhibition (36). Furthermore, responses to drug combinations were primarily assessed using short-term viability assays and fixed treatment conditions, which cannot fully recapitulate clinically relevant treatment durations and exposure patterns.

In conclusion, integrated functional and molecular analyses of resistance to BRAF-targeted therapy in preclinical models of CRC identified actionable vulnerabilities and highlighted convergent reactivation of MAPK signaling as a recurrent mechanism of therapeutic escape, frequently involving RAS alterations. This study provides proof-of-concept that combined BRAF and RAS inhibition can overcome acquired resistance driven by emerging RAS mutations and support further evaluation of this strategy in *BRAF*^V600E^-mutated CRC.

## Materials and Methods

### Cell line models and culture conditions

The human cancer cell lines COLO-205 (RRID:CVCL_0218) and HCT-116 (RRID:CVCL_0291) were acquired from the National Cancer Institute (NCI)-60 panel (Frederick, MD, USA), and the American Type Culture Collection (ATCC; Manassas, VA, USA), respectively (13). HT-29 (RRID:CVCL_0320) was kindly provided by R. Hamelin (National Institute for Health and Medical Research [INSERM], France). HCT-116 CRISPR-engineered variants harboring homozygous (catalog #SL731) and heterozygous (catalog #SL732) *BRAF*^V600E^ mutations were obtained from GeneCopoeia, Inc. (Rockville, MD, USA). Previously published in-house genetic, transcriptomic, and drug sensitivity data were included for additional CRC cell lines (13,14).

The generation of resistant sublines to encorafenib (HT-29^E^ and COLO-205^E^) and the combination of encorafenib and afatinib (HT-29^AE^, HT-29^EAE^, COLO-205^AE^, and COLO-205^EAE^) is detailed in the Results section (Fig. S3). Briefly, drug-resistant variants were generated by periodic treatment with gradually increasing concentrations of encorafenib (E) or the combination of encorafenib and afatinib (AE and EAE). The starting concentrations were the IC_50_ values for each cell line, and cells at approximately 80% confluency were exposed to the respective compound. After the emergence of drug-resistant colonies, cells were passaged and the concentration of the selection drug was increased 2-fold once cell proliferation remained undisturbed for at least two consecutive passages. Drug dosages were progressively increased until the cells became completely resistant to the highest concentration of the selection drug (Fig. S3). Samples were harvested at several timepoints, including totally 7 and 5 sublines of HT-29 and COLO-205, respectively.

Cell lines were maintained in antibiotic-free Dulbecco’s Modified Eagle Medium/Nutrient Mixture F-12 (DMEM/F-12 #11320033; Thermo Fisher Scientific, Waltham, MA, USA) supplemented with 2 mM GlutaMAX (#35050061; Thermo Fisher Scientific) and 15% fetal bovine serum (FBS #F7524; Sigma-Aldrich, St. Louis, MO, USA) at 37 °C in a humidified incubator with 5% CO₂. Mycoplasma contamination was routinely tested using the MycoAlert Mycoplasma Detection Assay (#LT07-710; Lonza, Basel, Switzerland). Parental cell lines and derived sublines were authenticated by short tandem repeat (STR) profiling using the AmpFLSTR Identifiler PCR Amplification Kit (#4427368; Thermo Fisher Scientific).

### Patient-derived tumor organoids

In the present study, 17 *BRAF*-mutated PDOs generated from liver metastases of patients with CRC have been analyzed (Table S1). The lineages are from patients enrolled in a prospective observational study of liver resection for metastatic CRC at Oslo University Hospital, and 13 of the *BRAF*^V600E^-mutated PDOs have recently been reported (15). All patient-derived material and associated clinical data were collected and used in accordance with approved ethical and data protection protocols. Written informed consent was obtained from all the patients, and the study was conducted in accordance with the Declaration of Helsinki. Approvals were granted by the Norwegian Data Protection Authority and the Regional Committee for Medical and Health Research Ethics, South-Eastern Norway (REC numbers 1.2005.1629, 2010/1805, and 2017/780).

Tumor specimens were mechanically dissociated into small cell clusters, filtered through a 70 µm strainer, embedded in growth factor-reduced Matrigel (Corning Inc., Corning, NY, USA), and cultured in Advanced DMEM/F-12 (#12634010; Thermo Fisher Scientific) supplemented with Y-27632, HEPES, GlutaMAX, penicillin-streptomycin, B27 supplement, [Leu15]-Gastrin I, N-acetyl-L-cysteine, EGF, Noggin, A83-01, and SB202190 to support organoid establishment and maintenance (15). PDOs were passaged every 7-10 days by enzymatic dissociation using TrypLE Express (#12604013; Thermo Fisher Scientific) and re-seeded in fresh Matrigel. Culture medium was replenished every 7 days. Mycoplasma contamination was routinely tested using the MycoAlert Mycoplasma Detection Assay (#LT07-710; Lonza). PDO identity was verified by STR profiling using the AmpFLSTR Identifiler PCR Amplification Kit (#4427368; Thermo Fisher Scientific) and was confirmed by comparison to the corresponding tumor tissue. Previously published PDOs have been characterized by targeted mutation profiling using a 20-gene panel, transcriptomic profiling, and sensitivity testing using panels comprising 41-47 compounds (15).

### Single-agent and combination drug screening

Previously generated high-throughput drug sensitivity data from cell lines (14) and PDOs (15), were reanalyzed in the present study with a focus on *BRAF*-mutated models.

Parental cell lines and derived sublines of HT-29, COLO-205, and HCT-116 (n = 12) were analyzed by medium-throughput drug sensitivity screening. The single-agent screening panel comprised 41 small-molecule inhibitors at eight concentrations and six drug combinations were tested at a 1:1 concentration ratio across seven concentrations (Table S4).

In addition, a customized screen with 22 unique drug combinations was designed for this study and performed across 13 models, including nine cell lines and four PDOs. Combination partners with BRAF inhibitors were selected based on transcriptomic analyses indicating pathway activation in resistant models (Table S5), and prior preclinical and clinical evidence of established pathway interactions and activity in CRC and other cancer types. All drug pairs were preprinted as full 7 × 7 concentration matrices in 384-well plates using acoustic dispensing in collaboration with the FIMM High Throughput Biomedicine Unit, which is hosted by the University of Helsinki and supported by HiLIFE and Biocenter Finland. Cell lines were seeded at a density of 2-4 × 10³ cells per well and PDOs at 500-600 structures per well (40 – 70 µm), in a final volume of 40 µL per well.

Combinations of encorafenib with the proteasome inhibitors bortezomib and carfilzomib, the selective SRC inhibitor eCF506, the panRAS inhibitors daraxonrasib and RMC-7977, or a SHOC2-RAS interaction inhibitor (PPI-IN-1) were tested in full dose-response matrices spanning 4-9 concentrations using manual drug dispensing (Table S6). For these experiments, drugs were added 24 h after cell seeding, in contrast to cells seeded into preprinted drug plates that were directly treated at the time of seeding.

Cells were exposed to single-agents or drug combinations for 72 h for cell lines and 96 h for PDOs, as previously described (15–17). Cell viability was assessed using the CellTiter-Glo 3D Cell Viability Assay (#G9681; Promega, Madison, WI, USA), and luminescence was measured using a Victor 3 microplate reader (PerkinElmer, Waltham, MA, USA). Raw luminescence signals were normalized to wells with benzethonium chloride as a positive control and DMSO as a negative control to calculate the percent viability. Dose response curves were fitted with the *drums* R package (v. 0.9.1; https://github.com/sveenlab/drums), incorporating logistic regression implemented with the *logLogisticRegression* function in the R package PharmacoGx (v. 3.0.2) and DSS estimation (18).

Drug interactions were quantified using SynergyFinder 3.0 (RRID:SCR_019318) (19). For each drug pair, maximum and mean percent viability values and maximum and mean synergy scores were calculated across the full combination matrix. Biological and technical replicates of combination treatments involving identical drug pairs or agents targeting similar pathways demonstrated high reproducibility, as reflected by the strong correlations between synergy scores (Fig. S16A-D). The Bliss independence model was selected for downstream analyses because its synergy scores showed the strongest correlations with those obtained using the Zero Interaction Potency (ZIP) and Highest Single Agent (HSA) models (Fig. S16E).

### DNA/RNA extraction, MSI status and gene panel sequencing

DNA and RNA extraction using the AllPrep DNA/RNA/miRNA Universal Kit (#80224, Qiagen, Hilden, Germany), determination of MSI status by analysis of the BAT-25/26 loci or by using the MSI Analysis System v1.2 (Promega, Madison, WI, USA), and targeted DNA sequencing of a custom CRC-relevant panel (n = 20 genes) have previously been published for an in-house collection of 103 parental cell lines and 150 PDOs (13,15). Resistant cell line models, CRISPR-modified cell lines and four additional *BRAF*^V600E^-mutated PDOs included in this study were analyzed by the same protocols (n = 16, Table S9). Sequencing was performed on the Illumina MiniSeq system in 2x73 base pair paired-end mode to a median coverage of 544X (range 281– 821). Processing of raw sequencing reads was done as previously described (13), including sequence alignment to the GRCh38 human reference genome using BWA (v0.7.17, RRID:SCR_010910), refinement of sequencing reads using Picard (v2.19.0, RRID:SCR_006525) and GATK (v4.1.2 for cell lines and v4.5.0 for PDOs, RRID:SCR_001876), mutation calling with Mutect2 in tumor-only mode, annotation by ANNOVAR (v2016-Feb-01, RRID:SCR_012821) and filtering of the candidate mutation list. DNA copy number aberrations were identified using the MiniCN (v0.1.0) R package (https://github.com/SveenLab/miniCN).

### Gene expression profiling and data analyses

Derivates of HT-29, COLO-205 and HCT-116 were analyzed for gene expression using the Affymetrix Human Transcriptome Array 2.0 (HTA; Themo Fisher Scientific) with 100 ng of total RNA as input and following the manufacturer’s protocol. The in-house collection of parental cell lines and a subset of PDOs have previously been analyzed on HTA (14,15,20). Raw intensity CEL-files were processed in three separate datasets (parental cell lines, *n* = 103; CRISPR-modified and resistant cell lines, *n* = 16; PDOs, *n* = 12) using the robust multi-array average method (21), implemented with the *justRMA* function in the R package *affy* (v1.80.0, RRID:SCR_012835) (22) and custom Brainarray CDF files (hta20hsentrezgcdf_25.0.0) (23). Gene annotations based on the GRCh38 genome assembly were retrieved using the *getBM* function in the R package *biomaRt* (v2.58.0, RRID:SCR_019214) (24), and only protein-coding genes with annotated HGNC symbols were retained. Principal component analysis was performed using the R package *FactoMineR* (v2.9, RRID:SCR_014602) on the 3,000 genes with the largest 10^th^-90^th^ percentile expression ranges across samples. Gene set enrichment analysis was performed using a custom gene set collection (*n* = 124 gene sets; Table S2). Group-wise comparisons were performed using the *GSA* function in the R package *GSA* (v.1.03.2) (25), whereas single-sample enrichment scores were calculated using the *gsva* function in the R package *GSVA* (v.1.50.0) (26).

### Immunoblotting

Parental and resistant cell lines (2.5-6.2 x10^5^ cells/well in 6-well plates) were treated for 72 h with vehicle control (0.01% DMSO), encorafenib, daraxonrasib, or RMC-7977, as single agents, or with combinations of encorafenib with daraxonrasib or RMC-7977. Each drug was used at a concentration of 1 µM. Following treatment, cells were collected, washed once with PBS, and lysed in 150 µl SDS electrophoresis sample buffer composed of 10 mM Tris (pH 6.8), 15% (w/v) glycerol, 3% (w/v) SDS, 0.01% (w/v) bromophenol blue, and 5% (v/v) 2-mercaptoethanol. Lysates were sonicated for 25 seconds at 37% amplitude and subsequently denatured at 95 °C for 5 minutes. Protein concentrations were determined using the RC DC Protein Assay kit (#5000119; Bio-Rad, Hercules, CA, USA) according to the manufacturer’s instructions.

Equal amounts of protein were resolved by SDS-PAGE on 10% polyacrylamide gels and transferred to PVDF membranes using the Trans-Blot Turbo Transfer System (Bio-Rad). Membranes were blocked and incubated overnight at 4 °C with primary antibodies against phospho-ERK1/2 (#9101; Cell Signaling, Danvers, MA, USA; 1:1,000; RRID:AB_331646), total ERK1/2 (#9102; Cell Signaling; 1:1,000; RRID:AB_330744), and β-actin (#A2228; Sigma-Aldrich; 1:5,000; RRID:AB_476697). After washing, total ERK1/2 and β-actin were detected using IRDye 800CW goat anti-rabbit IgG (#926-32211; LI-COR; 1:10,000; RRID:AB_621843) and IRDye 680RD goat anti-mouse IgG (#926-68070; LI-COR; 1:10,000; RRID:AB_10956588) respectively, diluted in Tris-buffered saline containing Tween 20 (TBS-T) and 5% milk. Phospho-ERK1/2 was detected using an HRP-conjugated secondary antibody (#1706515 Bio-Rad; 1:5,000) and SuperSignal West Dura Extended Duration Substrate kit (Thermo Fisher Scientific). Fluorescent and chemiluminescent signals were captured using the ChemiDoc™ MP imaging system (Bio-Rad), and band densitometry was performed using Image Lab software (v6.1; Bio-Rad).

### Phospho-kinase profiling

Phosphorylation profiles were assessed in parental HT-29 and resistant HT-29^AE^ cells using the Proteome Profiler Human Phospho-Kinase Array Kit (#ARY003C; R&D Systems, Minneapolis, MN, USA), which detects phosphorylation levels at specific sites of 37 proteins and total amounts of two proteins. Cells were treated with the combination of encorafenib and daraxonrasib, each at 1 µM, or DMSO for 72 h. Cell lysates were prepared and analyzed according to the manufacturer’s instructions. Protein concentration was determined with the RC DC Protein Assay kit and 350 µg of total protein was used per array set. Following blocking, membranes were incubated with cell lysates overnight, washed, and sequentially incubated with biotinylated detection antibodies and streptavidin-HRP. Signals were developed using the supplied chemiluminescent reagents, captured using the ChemiDoc™ MP imaging system (Bio-Rad) and quantified using Image Lab software (Bio-Rad). Background correction was performed using the designated negative-control PBS spots, and signal intensities were normalized to the reference-control spots on the corresponding membrane (Supplementary Table 10).

Normalized intensities of duplicate capture-antibody spots were analyzed as the mean for each target. Treatment responses were analyzed as log₂ fold changes relative to the corresponding DMSO control within each cell line and subsequently compared between the cell lines [Δlog₂FC = log₂FC(HT-29^AE^) − log₂FC(HT-29)]. Targets with duplicate-spot coefficients of variation >25% in any condition were retained and flagged with an asterisk.

### Statistical analyses

All statistical analyses and data visualizations were performed using R (v4.2.1) (27). Correlations were performed using the *cor.test* function with either the Spearman or Pearson method, as indicated. Two-sided Wilcoxon rank-sum and signed-rank tests were performed using the *wilcox_test* function in the *rstatix* package, as appropriate. For analyses involving multiple comparisons, p-values were adjusted using the Benjamini-Hochberg method. Heatmaps were generated using the *pheatmap* package (RRID:SCR_016418) with complete-linkage hierarchical clustering. Statistical significance was defined as p-value or adjusted p-value < 0.05.

## CRediT authorship contribution statement

**Kushtrim Kryeziu**: Conceptualization, Investigation, Data curation, Formal analysis, Funding acquisition, Writing – original draft. **Jakob M. Stenersen**: Investigation, Formal analysis. **Luís Nunes**: Formal analysis, Visualization, Writing – review & editing. **Max Z. Totland**: Investigation, Formal analysis. **Ida Sophie Gjøstøl Strømsvåg:** Validation, Writing – review & editing. **Nora S. Strømnes**: Investigation, Formal analysis. **Vilde C. W. Elster**: Investigation, Formal analysis. **Edward Leithe**: Writing – review & editing. **Barbara Niederdorfer**: Investigation, Validation. **Visame Jonuzi**: Investigation, Formal analysis. **Ina A. Eilertsen**: Data curation, Formal analysis. **Ragnhild A. Lothe**: Supervision, Resources, Funding acquisition, Writing – review & editing. **Anita Sveen**: Conceptualization, Investigation, Supervision, Formal analysis, Funding acquisition, Project administration, Writing – original draft.

## Acknowledgments

We thank Mette Eknæs and Merete Hektoen for their valuable technical assistance.

This work was supported by The Norwegian Cancer Society, project numbers 208336 and 246954 to AS, 297971 to KK, 223319 to RAL and by the South-Eastern Norway Regional Health Authority, project number 2023101 to AS, 2024059 to KK and 2024108 to RAL.

## Data availability

Previously published data on cell lines and PDOs reanalyzed in this study are available under (14,15). DSS for single drugs and percent viability inhibition for drug combinations are available in Supplementary Tables S7 and S8, respectively. Mutation data for the parental and resistant cell lines, CRISPR-modified cell lines and newly sequenced PDOs are available as Supplementary Table S9. Phospho-kinase array data are available in Supplementary Table S10. Gene expression data are available in Gene Expression Omnibus (GEO, RRID:SCR_005012) repository under accession GSE97023, GSE309172 and GSE294511. The gene expression data from parental and derivates of HT-29, COLO-205 and HCT-116 have been deposited to GEO under accession GSE350004. All data were processed and analyzed using published software or code referenced in this study, and no custom code was developed.

