## Supplementary Figures for "Combined BRAF and RAS inhibition overcomes acquired resistance in *BRAF*-mutated colorectal cancer"

Address:

**Running Title:** RAS and BRAF inhibition targets resistant colorectal cancer

**Keywords:** CRC, BRAF, RAS, drug resistance, cell line, patient-derived organoid, pharmacogenomics.

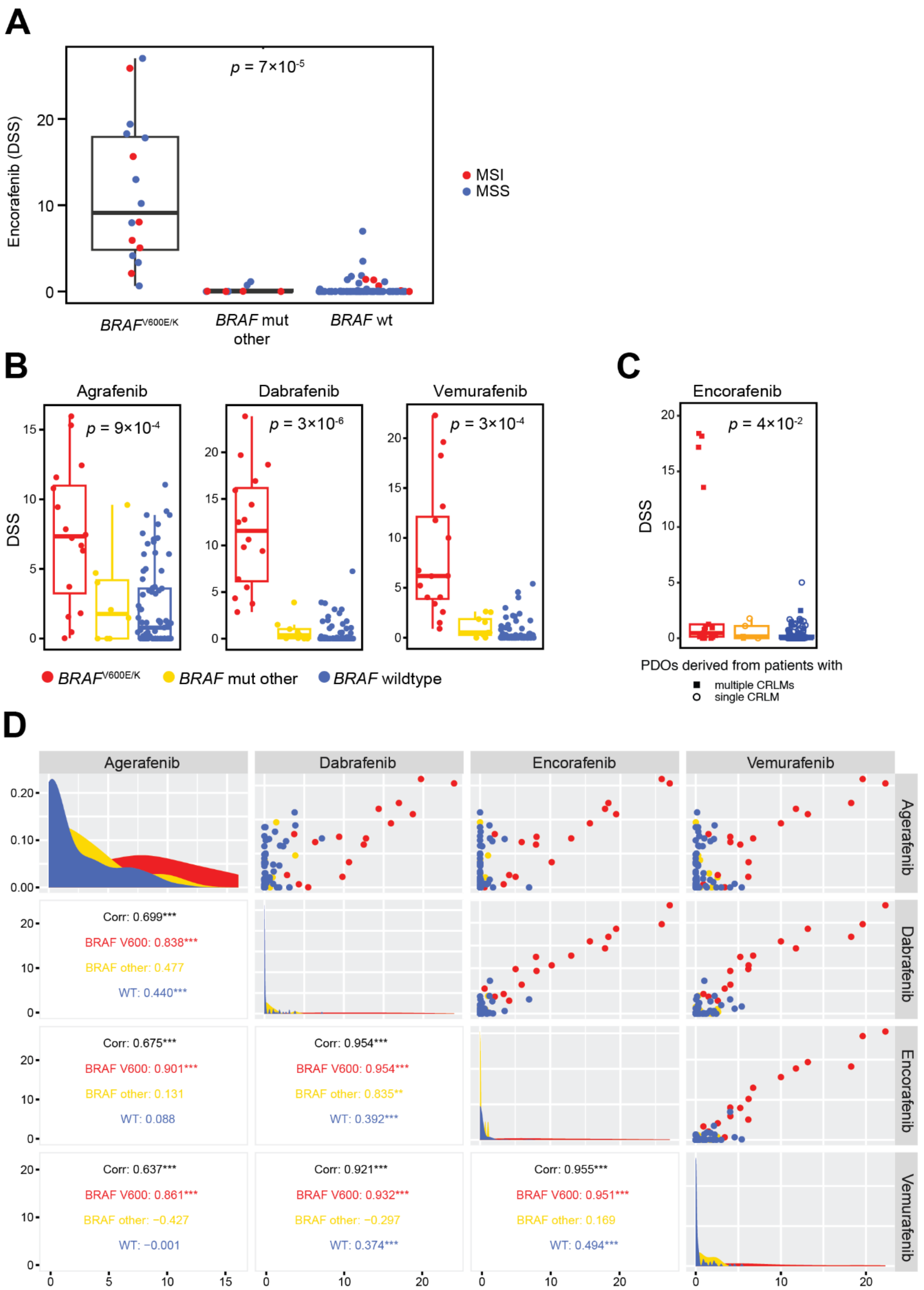


**Supplementary Figure 1. Molecular and BRAF inhibitor sensitivity patterns for *BRAF*$^{V600E/K}$-mutated cell lines and PDOs.**

**A**) Box plot of DSS to single agent encorafenib according to *BRAF* mutation status and microsatellite stability status in CRC cell lines (total $n$ = 102). DSS distributions for agerafenib, dabrafenib, encorafenib, and vemurafenib across cell lines (**B**) and encorafenib in PDOs (**C**) grouped by *BRAF* mutation status. Statistical comparisons between *BRAF*$^{V600E}$ and all other samples were performed using Welch's t-test. Filled symbols in panel **C** indicate PDOs derived from patients with multiple colorectal liver metastases (CRLMs). The PDO analysis was based on previously published drug-sensitivity data (15) and was reanalyzed here after inclusion of four additional PDOs that were molecularly characterized in the present study and classified as *BRAF*$^{V600E}$-mutated. **D**) Pairwise correlation analysis of DSS values for agerafenib, dabrafenib, encorafenib, and vemurafenib across cell lines. Density plots and scatter plots show overall correlations and correlations stratified by *BRAF* mutation status, with Pearson correlation coefficients and significance levels indicated.

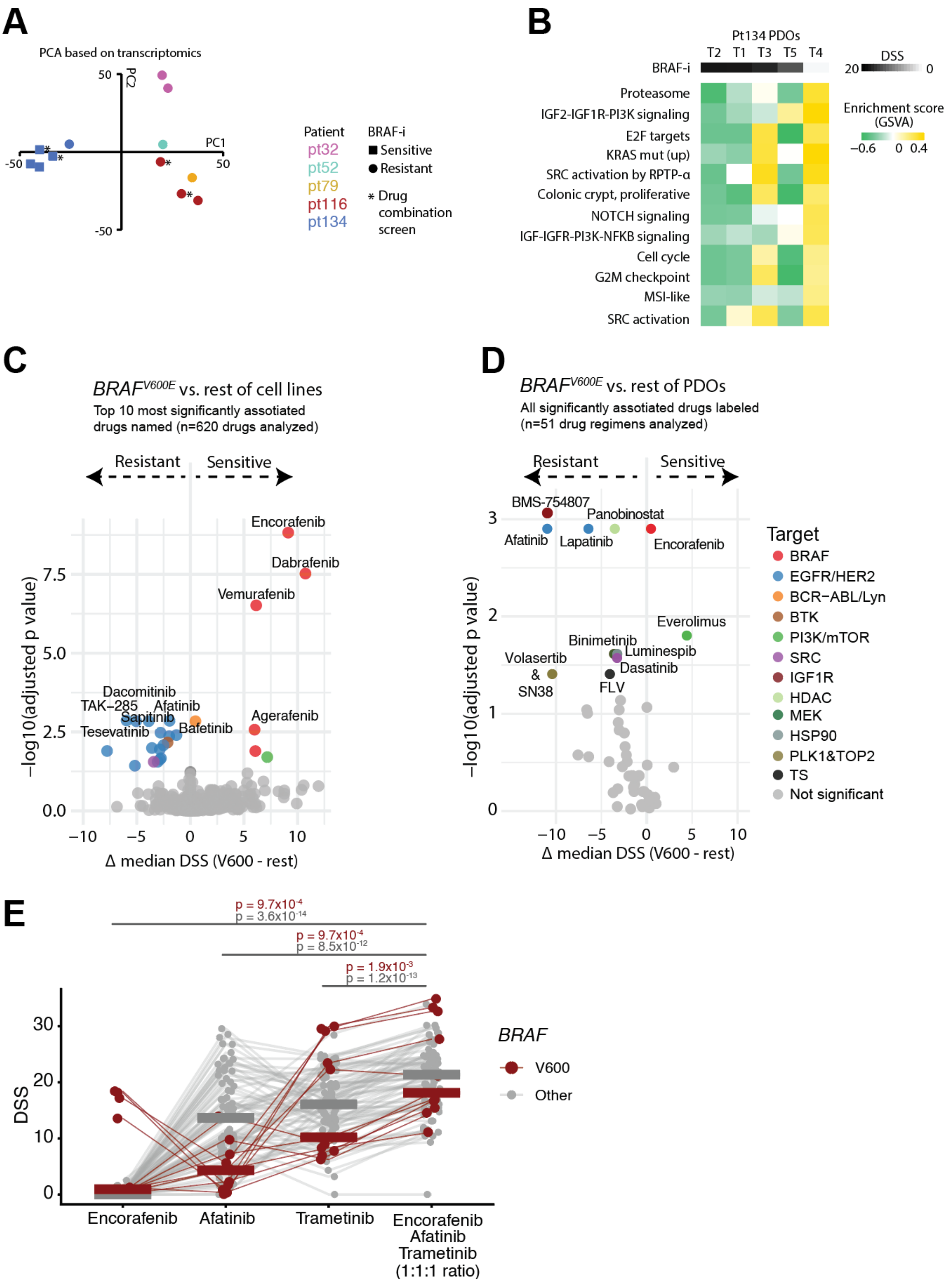
A
PCA based on transcriptomics
PC2
PC1
Patient
pt32
pt52
pt79
pt116
pt134
BRAF-i
Sensitive
Resistant
* Drug combination screen
B
Pt134 PDOs
T2 T1 T3 T5 T4
BRAF-i
DSS
Proteasome
IGF2-IGF1R-PI3K signaling
E2F targets
KRAS mut (up)
SRC activation by RPTP-α
Colonic crypt, proliferative
NOTCH signaling
IGF-IGFR-PI3K-NFKB signaling
Cell cycle
G2M checkpoint
MSI-like
SRC activation
Enrichment score (GSVA)
C
BRAF^V600E vs. rest of cell lines
Top 10 most significantly assotiated drugs named (n=620 drugs analyzed)
Resistant
Sensitive
Encorafenib
Dabrafenib
Vemurafenib
Dacomitinib
TAK−285
Afatinib
Sapitinib
Tesevatinib
Bafetinib
Agerafenib
−log10(adjusted p value)
Δ median DSS (V600 - rest)
D
BRAF^V600E vs. rest of PDOs
All significantly assotiated drugs labeled (n=51 drug regimens analyzed)
Resistant
Sensitive
BMS-754807
Panobinostat
Afatinib
Lapatinib
Encorafenib
Everolimus
Binimetinib
Luminespib
Volasertib & SN38
Dasatinib
FLV
−log10(adjusted p value)
Δ median DSS (V600 - rest)
Target
BRAF
EGFR/HER2
BCR−ABL/Lyn
BTK
PI3K/mTOR
SRC
IGF1R
HDAC
MEK
HSP90
PLK1&TOP2
TS
Not significant
E
p = 9.7x10^-4
p = 3.6x10^-14
p = 9.7x10^-4
p = 8.5x10^-12
p = 1.9x10^-3
p = 1.2x10^-13
DSS
Encorafenib
Afatinib
Trametinib
Encorafenib Afatinib Trametinib (1:1:1 ratio)
BRAF
V600
Other

**Supplementary Figure 2. Transcriptomic heterogeneity and association of *BRAF*$^{V600E}$-mutation with drug sensitivity in cell lines and PDOs.**

**A**) Principal components analyses based on genes (*n* = 3,000) with highest 10-90$^{th}$ percentile expression ranges among *BRAF*$^{V600E}$-mutated PDOs (*n* = 12 PDOs from 5 patients). Asterisks indicate PDOs included in drug combination screening. **B**) Single-sample enrichment scores (GSVA) of the top 12 gene sets ranked based on difference between an encorafenib resistant PDO and four sensitive PDOs from the same patient. DSS for encorafenib are indicated in the upper panel. Volcano plots highlighting the top 10 drugs most significantly associated with differential sensitivity between *BRAF*$^{V600}$-mutant and non-*BRAF*$^{V600}$ cell lines (n = 620 drugs) (**C**) and all drugs significantly associated in PDOs (n = 51 drugs regimens analyzed) (**D**). Drugs are colored according to their primary targets. **E**) Activity of encorafenib, afatinib, trametinib and their combination in 1:1:1 concentration ratio in *BRAF*-mutated PDOs (red) as compared to PDOs with non-V600 mutations and wild type status (grey). Horizontal lines of the respective groups indicate medians. P values are from paired Wilcoxon tests for the respective groups (*BRAF*$^{V600E}$-mutated models in dark red and grey models with wild type *BRAF* or non-V600E *BRAF* mutations) and treatments.

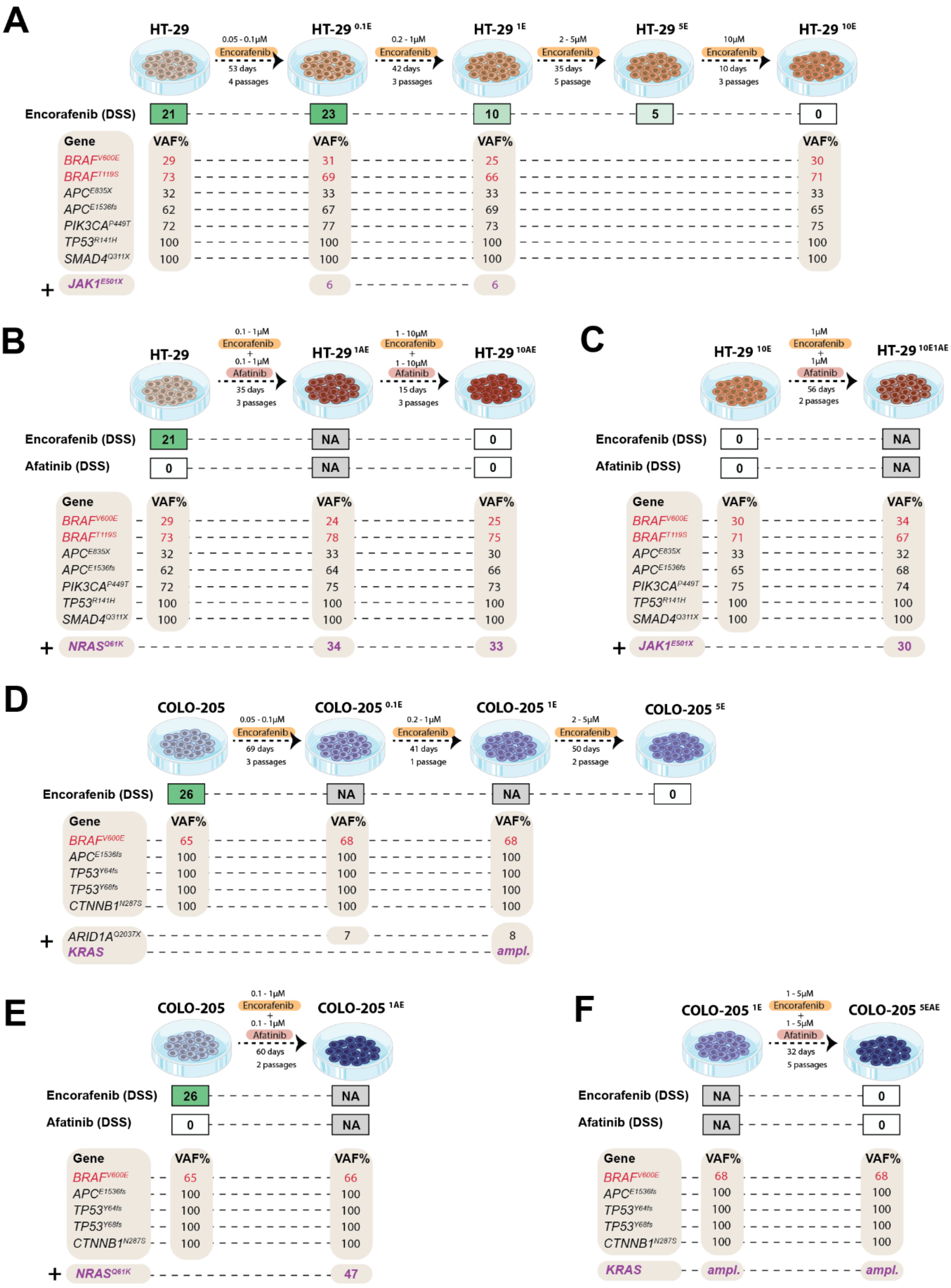
A
HT-29
0.05 - 0.1µM Encorafenib 53 days 4 passages
HT-29 0.1E
0.2 - 1µM Encorafenib 42 days 3 passages
HT-29 1E
2 - 5µM Encorafenib 35 days 5 passage
HT-29 5E
10µM Encorafenib 10 days 3 passages
HT-29 10E
Encorafenib (DSS) 21 23 10 5 0
Gene
VAF%
BRAF V600E 29 31 25 30
BRAF T119S 73 69 66 71
APC E835X 32 33 33 33
APC E1536fs 62 67 69 65
PIK3CA P449T 72 77 73 75
TP53 R141H 100 100 100 100
SMAD4 Q311X 100 100 100 100
+ JAK1 E501X 6 6
B
HT-29
0.1 - 1µM Encorafenib + 0.1 - 1µM Afatinib 35 days 3 passages
HT-29 1AE
1 - 10µM Encorafenib + 1 - 10µM Afatinib 15 days 3 passages
HT-29 10AE
Encorafenib (DSS) 21 NA 0
Afatinib (DSS) 0 NA 0
Gene
VAF%
BRAF V600E 29 24 25
BRAF T119S 73 78 75
APC E835X 32 33 30
APC E1536fs 62 64 66
PIK3CA P449T 72 75 73
TP53 R141H 100 100 100
SMAD4 Q311X 100 100 100
+ NRAS Q61K 34 33
C
HT-29 10E
1µM Encorafenib + 1µM Afatinib 56 days 2 passages
HT-29 10E1AE
Encorafenib (DSS) 0 NA
Afatinib (DSS) 0 NA
Gene
VAF%
BRAF V600E 30 34
BRAF T119S 71 67
APC E835X 33 32
APC E1536fs 65 68
PIK3CA P449T 75 74
TP53 R141H 100 100
SMAD4 Q311X 100 100
+ JAK1 E501X 30
D
COLO-205
0.05 - 0.1µM Encorafenib 69 days 3 passages
COLO-205 0.1E
0.2 - 1µM Encorafenib 41 days 1 passage
COLO-205 1E
2 - 5µM Encorafenib 50 days 2 passage
COLO-205 5E
Encorafenib (DSS) 26 NA NA 0
Gene
VAF%
BRAF V600E 65 68 68
APC E1536fs 100 100 100
TP53 Y64fs 100 100 100
TP53 Y68fs 100 100 100
CTNNB1 N287S 100 100 100
+ ARID1A Q2037X 7 8
KRAS ampl.
E
COLO-205
0.1 - 1µM Encorafenib + 0.1 - 1µM Afatinib 60 days 2 passages
COLO-205 1AE
Encorafenib (DSS) 26 NA
Afatinib (DSS) 0 NA
Gene
VAF%
BRAF V600E 65 66
APC E1536fs 100 100
TP53 Y64fs 100 100
TP53 Y68fs 100 100
CTNNB1 N287S 100 100
+ NRAS Q61K 47
F
COLO-205 1E
1 - 5µM Encorafenib + 1 - 5µM Afatinib 32 days 5 passages
COLO-205 5EAE
Encorafenib (DSS) NA 0
Afatinib (DSS) NA 0
Gene
VAF%
BRAF V600E 68 68
APC E1536fs 100 100
TP53 Y64fs 100 100
TP53 Y68fs 100 100
CTNNB1 N287S 100 100
KRAS ampl. ampl.

**Supplementary Figure 3. Stepwise acquisition of resistance to BRAF and EGFR inhibition in HT-29 and COLO-205 cell line models.**

**A)** Schematic overview of the stepwise generation of HT-29 sublines with increasing resistance to encorafenib through prolonged drug exposure at escalating concentrations. DSS for encorafenib and variant allele frequencies (VAFs) for selected genes are shown at each stage. Emergence and then disappearance of a truncating $\mathit{JAK1}^{E501X}$ mutation is indicated. **B)** Generation of HT-29 sublines with acquired resistance to combined encorafenib and afatinib treatment following prolonged dual-drug exposure, with corresponding DSS values and VAFs. Acquisition of an activating $\mathit{NRAS}^{Q61K}$ mutation is indicated. **C)** Exposure of encorafenib-resistant HT-29 cells to combined encorafenib and afatinib, with corresponding DSS values and emergence of $\mathit{JAK1}^{E501X}$. **D)** Stepwise generation of COLO-205 sublines with acquired resistance to encorafenib, showing encorafenib DSS values and VAFs for key driver genes. Acquisition of *KRAS* amplification and an $\mathit{ARID1A}^{Q2037X}$ mutation is indicated. **E)** Generation of COLO-205 sublines under combined encorafenib and afatinib exposure, with corresponding DSS values and emergence of $\mathit{NRAS}^{Q61K}$. **F)** Dual inhibition of encorafenib-resistant COLO-205 cells with encorafenib and afatinib, showing complete resistance and persistence of *KRAS* amplification. Drug concentrations, treatment durations, and passage numbers are indicated for each selection step. Numbers before the abbreviated encorafenib (E) and afatinib-encorafenib (AE) indicate the concentration of the respective drug or drug combination used for selection of resistance.

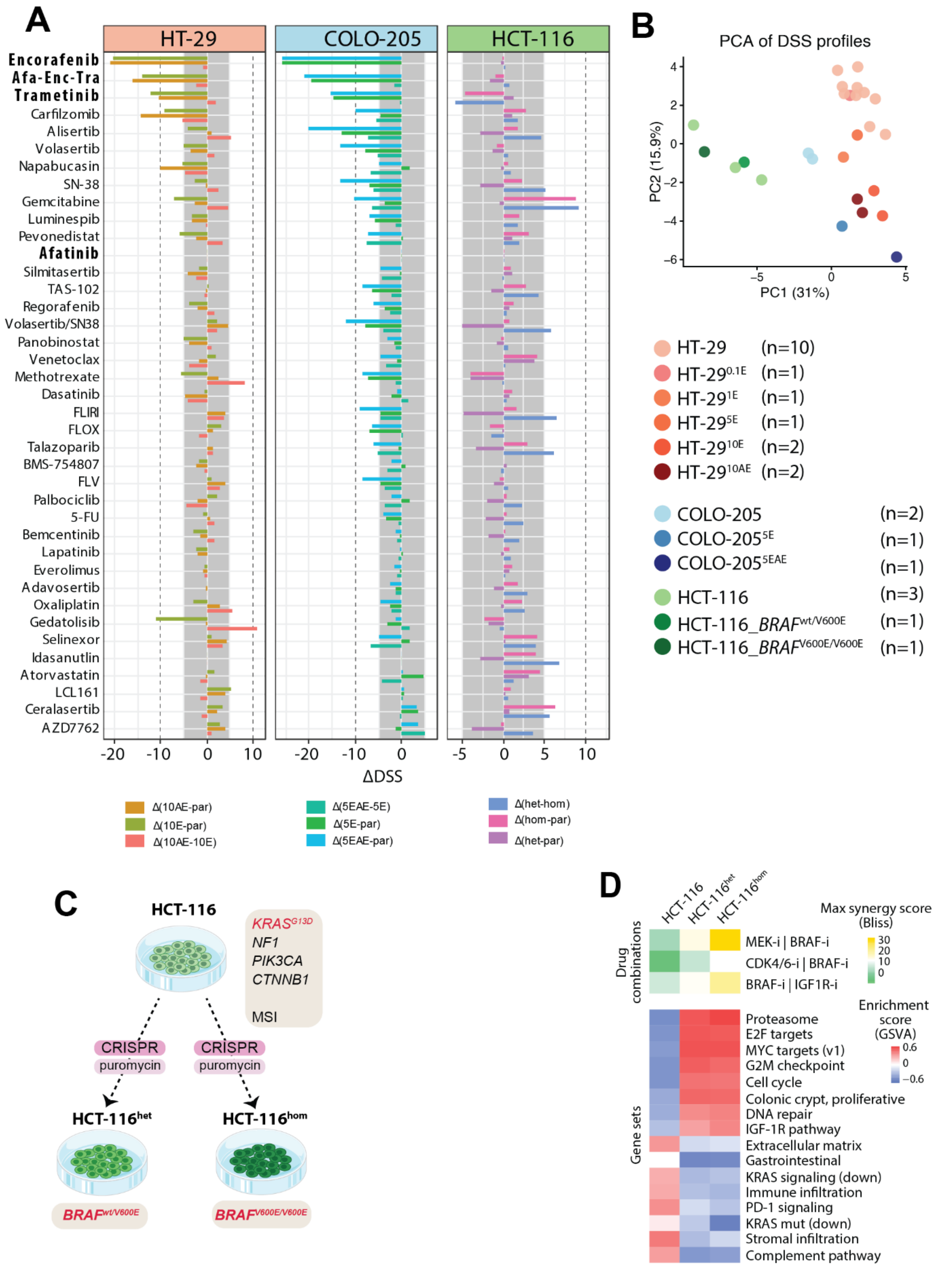
A
HT-29
COLO-205
HCT-116
Encorafenib
Afa-Enc-Tra
Trametinib
Carfilzomib
Alisertib
Volasertib
Napabucasin
SN-38
Gemcitabine
Luminespib
Pevonedistat
Afatinib
Silmitasertib
TAS-102
Regorafenib
Volasertib/SN38
Panobinostat
Venetoclax
Methotrexate
Dasatinib
FLIRI
FLOX
Talazoparib
BMS-754807
FLV
Palbociclib
5-FU
Bemcentinib
Lapatinib
Everolimus
Adavosertib
Oxaliplatin
Gedatolisib
Selinexor
Idasanutlin
Atorvastatin
LCL161
Ceralasertib
AZD7762
ΔDSS
Δ(10AE-par)
Δ(10E-par)
Δ(10AE-10E)
Δ(5EAE-5E)
Δ(5E-par)
Δ(5EAE-par)
Δ(het-hom)
Δ(hom-par)
Δ(het-par)
B
PCA of DSS profiles
PC2 (15.9%)
PC1 (31%)
HT-29 (n=10)
HT-29^0.1E (n=1)
HT-29^1E (n=1)
HT-29^5E (n=1)
HT-29^10E (n=2)
HT-29^10AE (n=2)
COLO-205 (n=2)
COLO-205^5E (n=1)
COLO-205^5EAE (n=1)
HCT-116 (n=3)
HCT-116_BRAF^wt/V600E (n=1)
HCT-116_BRAF^V600E/V600E (n=1)
C
HCT-116
KRAS^G13D
NF1
PIK3CA
CTNNB1
MSI
CRISPR
puromycin
CRISPR
puromycin
HCT-116^het
HCT-116^hom
BRAF^wt/V600E
BRAF^V600E/V600E
D
HCT-116
HCT-116^het
HCT-116^hom
Drug combinations
Max synergy score (Bliss)
MEK-i | BRAF-i
CDK4/6-i | BRAF-i
BRAF-i | IGF1R-i
Enrichment score (GSVA)
Gene sets
Proteasome
E2F targets
MYC targets (v1)
G2M checkpoint
Cell cycle
Colonic crypt, proliferative
DNA repair
IGF-1R pathway
Extracellular matrix
Gastrointestinal
KRAS signaling (down)
Immune infiltration
PD-1 signaling
KRAS mut (down)
Stromal infiltration
Complement pathway

**Supplementary Figure 4. Global drug response patterns and pathway dependencies across colorectal cancer cell line models.**

**A)** Changes in drug sensitivity (ΔDSS) relative to parental controls, drug-selection and edited variants for HT-29, COLO-205, and HCT-116 models, respectively. Positive values indicating increased sensitivity and negative values indicating increased resistance. **B)** PCA of DSS from 47 drugs across all tested cell line models and corresponding sublines, shown as pairwise comparisons. Samples are colored according to cell line identity and drug-selection, including parental and resistant HT-29 and COLO-205 sublines, as well as isogenic HCT-116 models. **C**) HCT-116 cell line and isogenic CRISPR–Cas9 engineered sublines with heterozygous HCT-116$^{het}$ or homozygous HCT-116$^{hom}$ *BRAF*$^{V600E}$ mutations. HCT-116 and the sublines harbor *KRAS*$^{G13D}$ and are microsatellite instable (MSI). CRISPR-edited cell lines were selected using puromycin. **D)** Heatmap summarizing maximum Bliss synergy scores for selected drug combinations (top 3 based on differential synergy scores) and corresponding top 16 gene sets based on differential single-sample enrichment scores (red: positive and blue: negative) between the parental, heterozygous, and homozygous *BRAF*$^{V600}$ models.

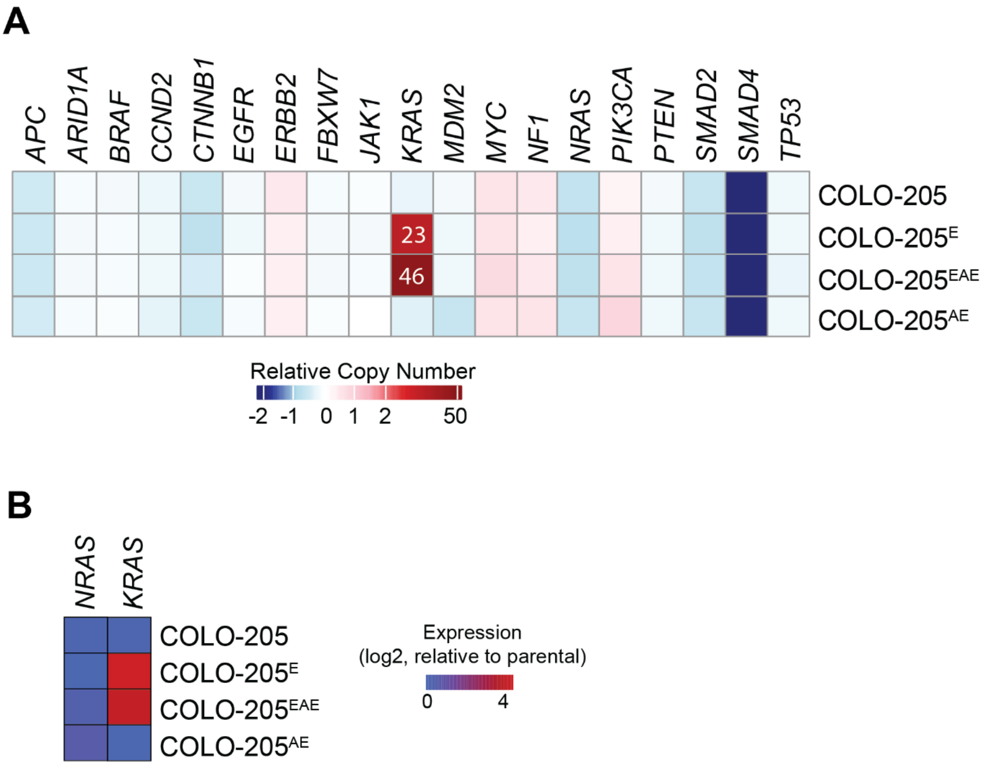


**Supplementary Figure 5. *KRAS* amplification and transcriptional increase of mRNA is associated with resistance to BRAF and EGFR inhibition in COLO-205.**

**A**) Heatmap of copy number alterations across a 20-gene panel in COLO-205 parental and resistant sublines. Color intensity reflects copy number relative to diploid (2 copies); blue indicates loss (minimum −2 as homozygous deletion) and red indicates amplification. **B**) *NRAS* and *KRAS* gene expression levels (log2 of the normalized mRNA expression) in COLO-205 parental cells and resistant sublines.

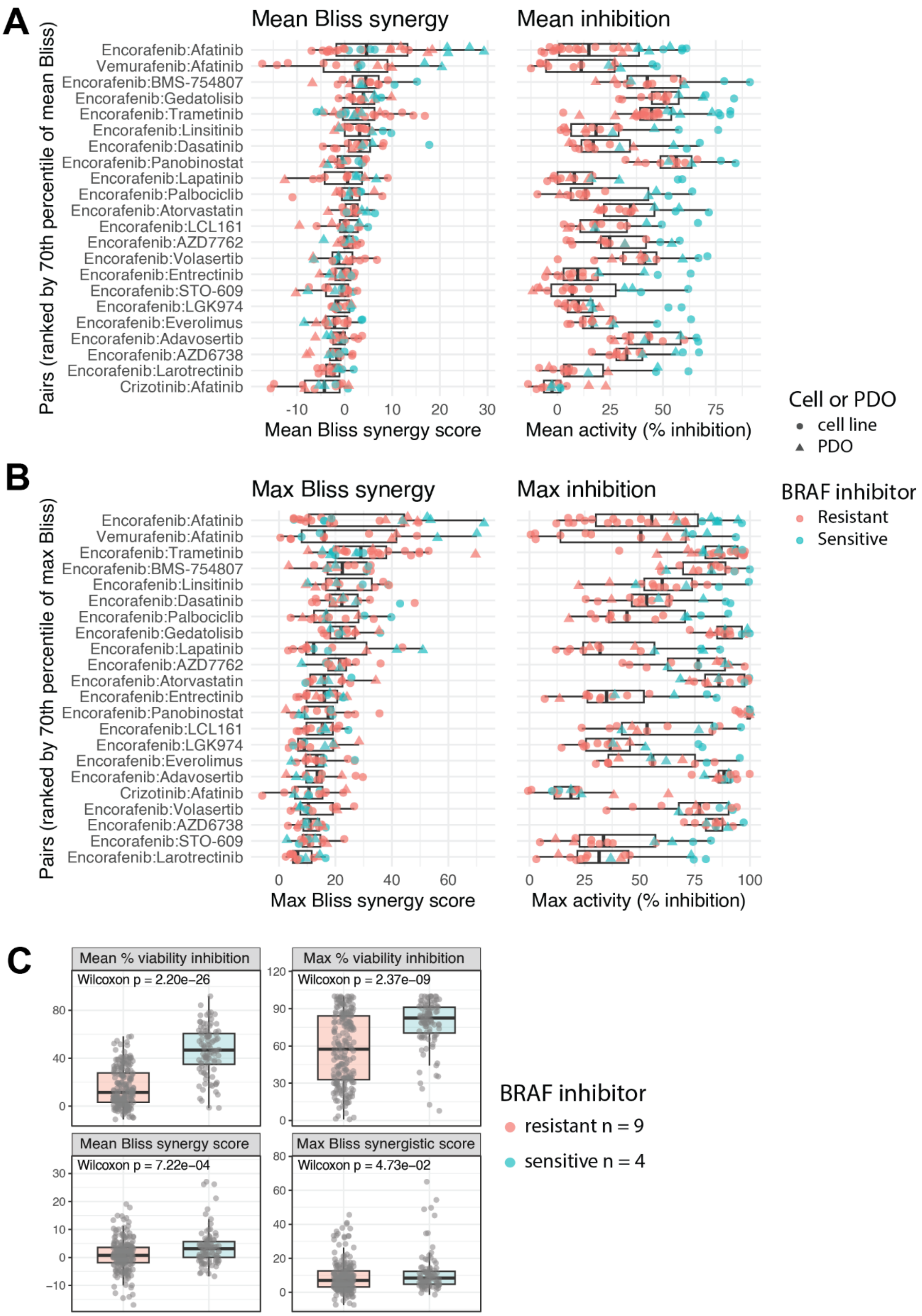
A
Mean Bliss synergy
Mean inhibition
Pairs (ranked by 70th percentile of mean Bliss)
Encorafenib:Afatinib
Vemurafenib:Afatinib
Encorafenib:BMS-754807
Encorafenib:Gedatolisib
Encorafenib:Trametinib
Encorafenib:Linsitinib
Encorafenib:Dasatinib
Encorafenib:Panobinostat
Encorafenib:Lapatinib
Encorafenib:Palbociclib
Encorafenib:Atorvastatin
Encorafenib:LCL161
Encorafenib:AZD7762
Encorafenib:Volasertib
Encorafenib:Entrectinib
Encorafenib:STO-609
Encorafenib:LGK974
Encorafenib:Everolimus
Encorafenib:Adavosertib
Encorafenib:AZD6738
Encorafenib:Larotrectinib
Crizotinib:Afatinib
Mean Bliss synergy score
Mean activity (% inhibition)
Cell or PDO
cell line
PDO
B
Max Bliss synergy
Max inhibition
Pairs (ranked by 70th percentile of max Bliss)
Encorafenib:Afatinib
Vemurafenib:Afatinib
Encorafenib:Trametinib
Encorafenib:BMS-754807
Encorafenib:Linsitinib
Encorafenib:Dasatinib
Encorafenib:Palbociclib
Encorafenib:Gedatolisib
Encorafenib:Lapatinib
Encorafenib:AZD7762
Encorafenib:Atorvastatin
Encorafenib:Entrectinib
Encorafenib:Panobinostat
Encorafenib:LCL161
Encorafenib:LGK974
Encorafenib:Everolimus
Encorafenib:Adavosertib
Crizotinib:Afatinib
Encorafenib:Volasertib
Encorafenib:AZD6738
Encorafenib:STO-609
Encorafenib:Larotrectinib
Max Bliss synergy score
Max activity (% inhibition)
BRAF inhibitor
Resistant
Sensitive
C
Mean % viability inhibition
Wilcoxon p = 2.20e−26
Max % viability inhibition
Wilcoxon p = 2.37e−09
Mean Bliss synergy score
Wilcoxon p = 7.22e−04
Max Bliss synergistic score
Wilcoxon p = 4.73e−02
BRAF inhibitor
resistant n = 9
sensitive n = 4

**Supplementary Figure 6. Drug combination efficacy and synergy stratified by sensitivity to BRAF inhibition across cell lines and patient-derived organoids.**

**A)** Mean Bliss synergy scores (left) and mean activity (% viability inhibition, right) as well as **B**) Maximum Bliss synergy scores (left) and maximum activity (% viability inhibition, right) for drug combinations tested in cell lines (n=9) and PDOs (n=4). Synergy scores are ranked based on the 70$^{th}$ percentile of mean synergy scores (**A**) or max synergy scores (**B**) for the indicated combinations. Each data point represents an individual model classified as sensitive (turquoise) or resistant (red) to encorafenib single treatment (based on DSS 5). **C)** Boxplots summarizing mean and maximum percent viability inhibition and mean and maximum Bliss synergy scores across all combinations in four BRAF inhibitor–sensitive and nine resistant models. Statistical comparisons were performed using the Wilcoxon test.

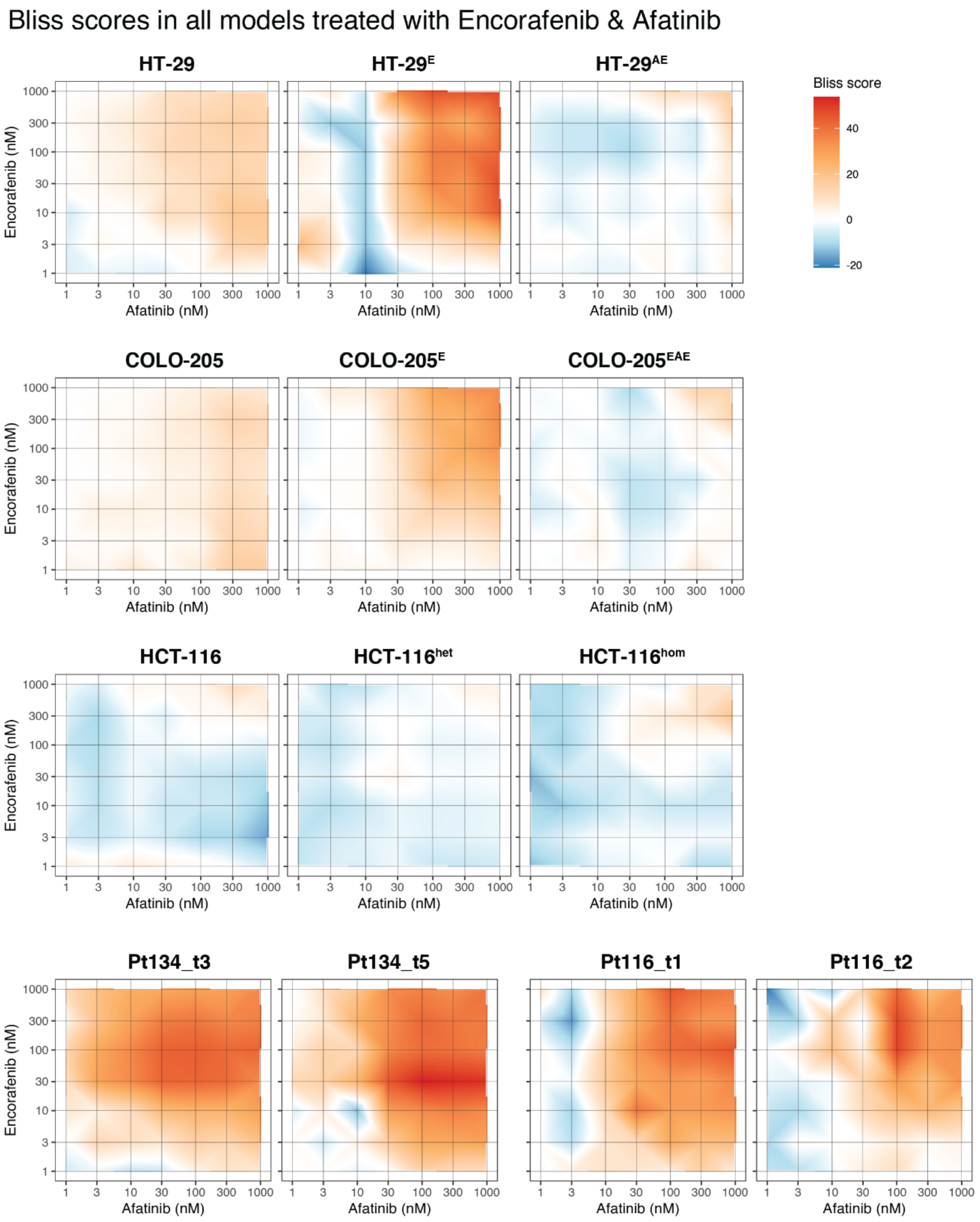


**Supplementary Figure 7. Bliss synergy landscapes for combined BRAF and EGFR inhibition across cell line and PDOs.**

Dose-response heatmaps showing Bliss scores for the combination of encorafenib and afatinib across parental and resistant HT-29 and COLO-205 sublines, isogenic HCT116 models, and selected PDOs. Bliss scores represent deviations from Bliss independence, with positive values indicating synergistic interactions and negative values indicating antagonism. Bliss score surfaces were interpolated (akima::interp, R) on log10 concentrations for visualization.

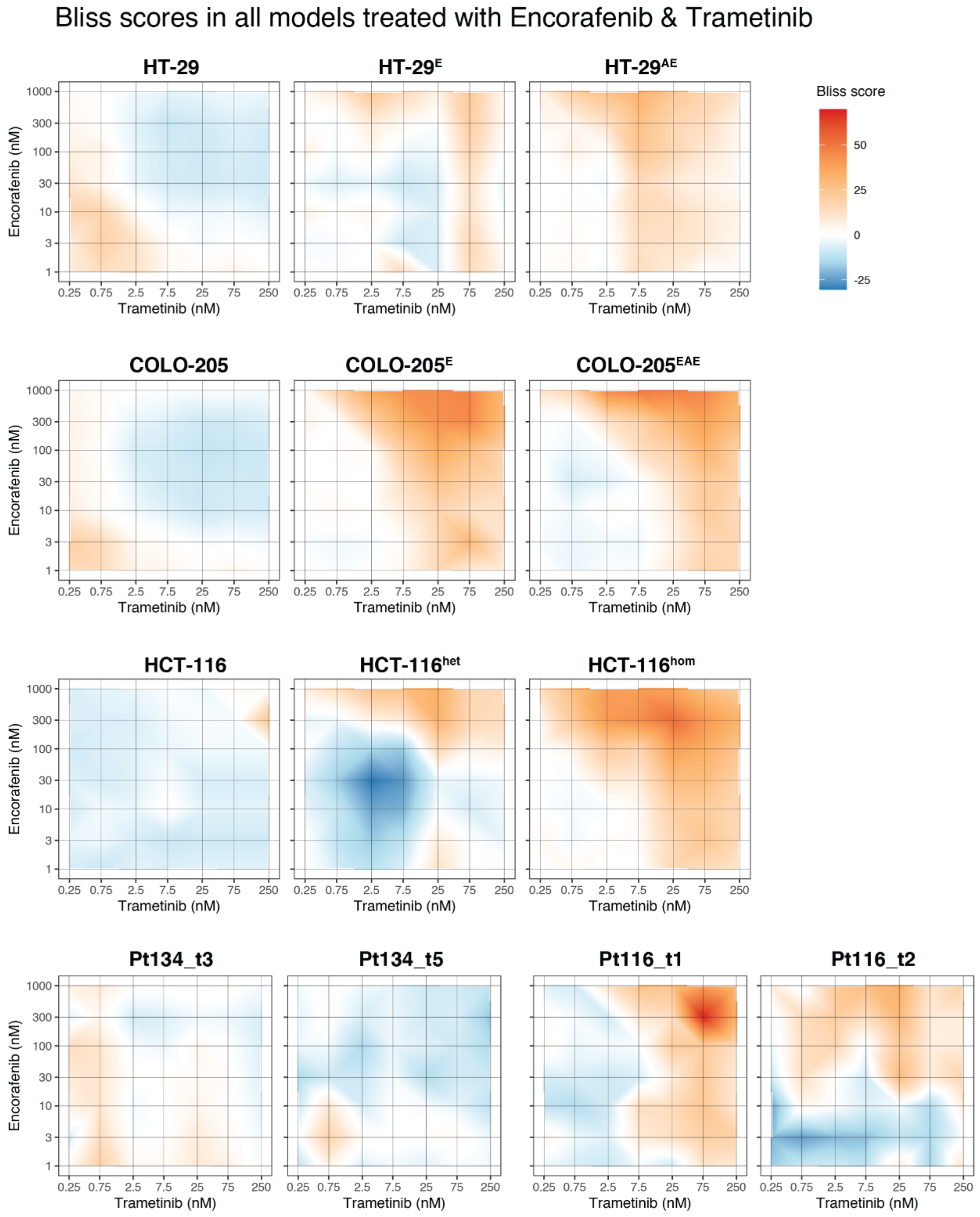


**Supplementary Figure 8. Bliss synergy landscapes for combined BRAF inhibitor encorafenib and MEK1/2 inhibitor trametinib across cell line and PDOs.**

Dose-response heatmaps showing Bliss scores for the combination of encorafenib and trametinib across parental and resistant HT-29 and COLO-205 sublines, isogenic HCT116 models, and selected PDOs. Bliss scores represent deviations from Bliss independence, with positive values indicating synergistic interactions and negative values indicating antagonism. Bliss score surfaces were interpolated (akima::interp, R) on log10 concentrations for visualization.

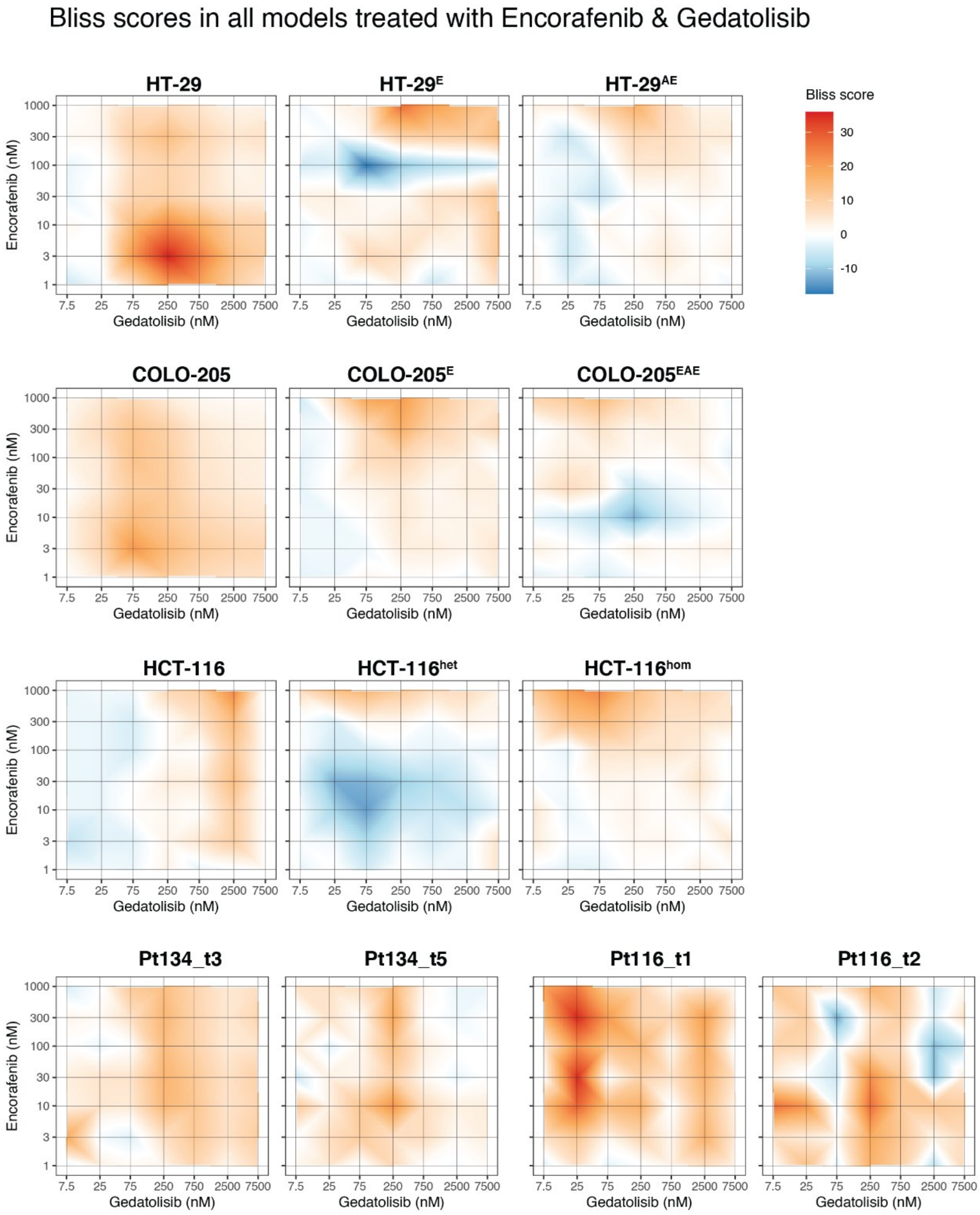


**Supplementary Figure 9. Bliss synergy landscapes for combined BRAF and PIK3A/mTOR inhibitor gedatolisib across cell line and PDOs.**

Dose-response heatmaps showing Bliss scores for the combination of encorafenib and gedatolisib across parental and resistant HT-29 and COLO-205 sublines, isogenic HCT116 models, and selected PDOs. Bliss scores represent deviations from Bliss independence, with positive values indicating synergistic interactions and negative values indicating antagonism. Bliss score surfaces were interpolated (akima::interp, R) on log10 concentrations for visualization.

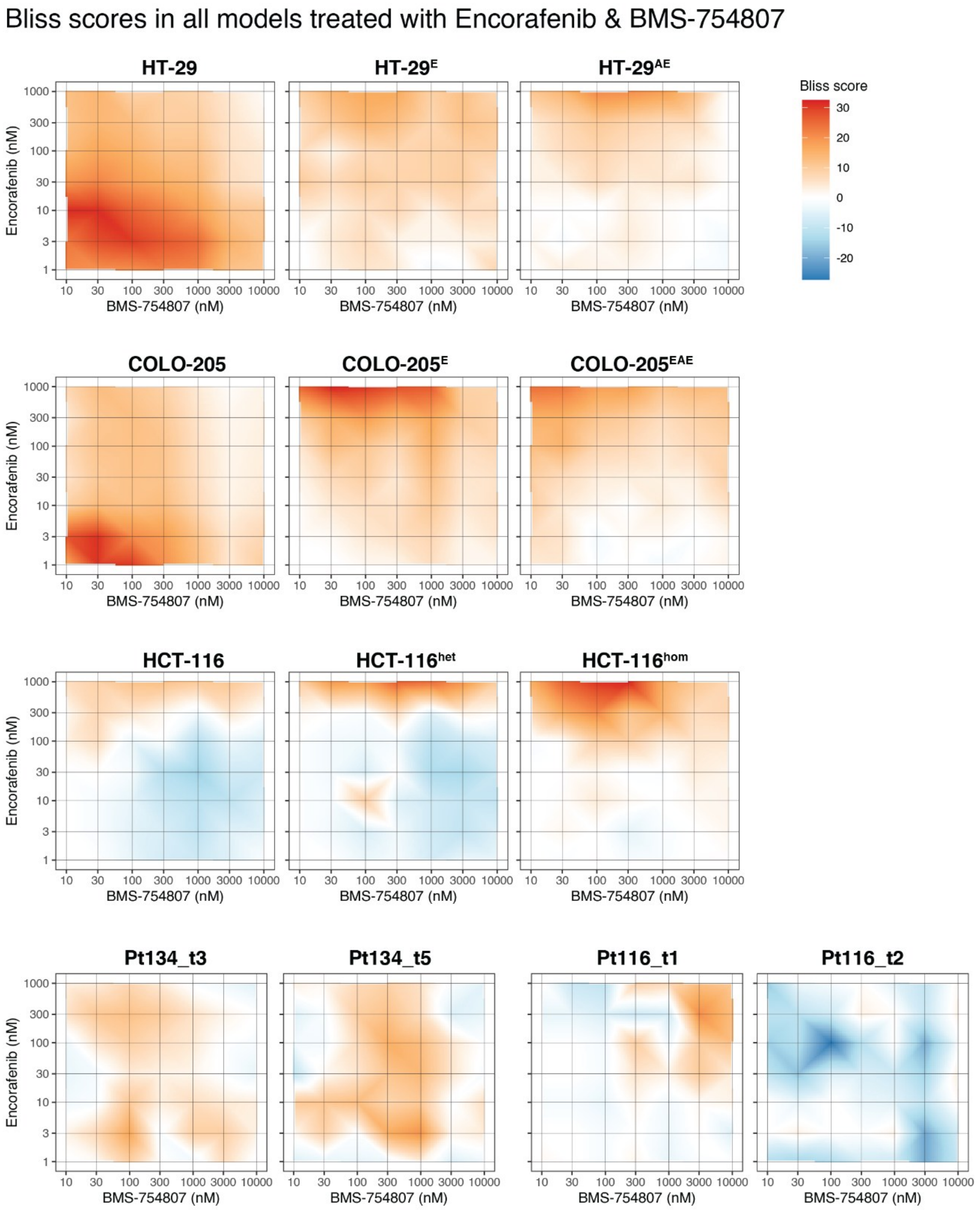


**Supplementary Figure 10. Bliss synergy landscapes for combined BRAF and IGF1R inhibitor BMS-754807 across cell line and PDOs.**

Dose-response heatmaps showing Bliss scores for the combination of encorafenib and BMS-754807 across parental and resistant HT-29 and COLO-205 sublines, isogenic HCT116 models, and selected PDOs. Bliss scores represent deviations from Bliss independence, with positive values indicating synergistic interactions and negative values indicating antagonism. Bliss score surfaces were interpolated (akima::interp, R) on log10 concentrations for visualization.

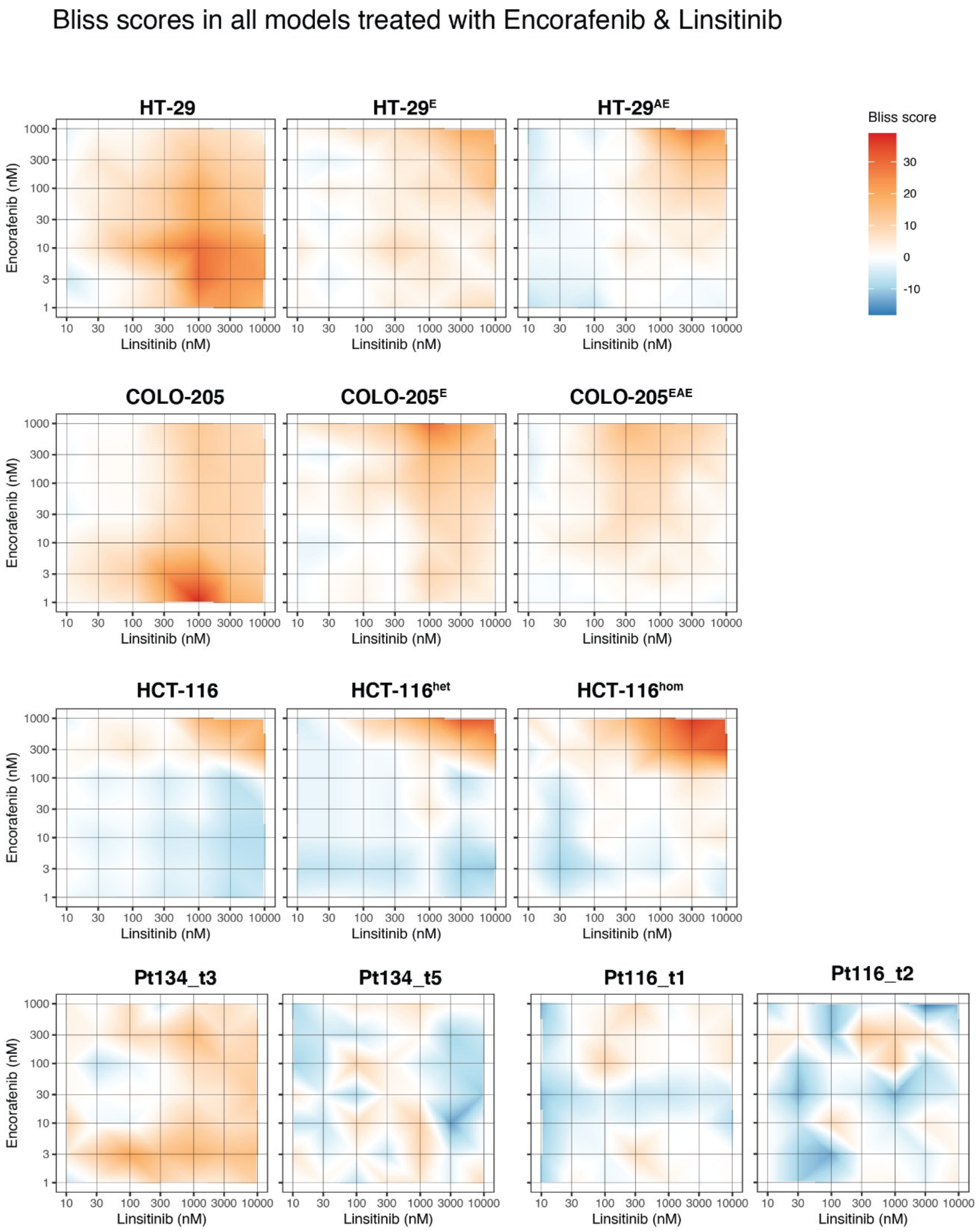


**Supplementary Figure 11. Bliss synergy landscapes for combined BRAF and IGF1R inhibitor linsitinib across cell line and PDOs.**

Dose-response heatmaps showing Bliss scores for the combination of encorafenib and linsitinib across parental and resistant HT-29 and COLO-205 sublines, isogenic HCT116 models, and selected PDOs. Bliss scores represent deviations from Bliss independence, with positive values indicating synergistic interactions and negative values indicating antagonism. Bliss score surfaces were interpolated (akima::interp, R) on log10 concentrations for visualization.

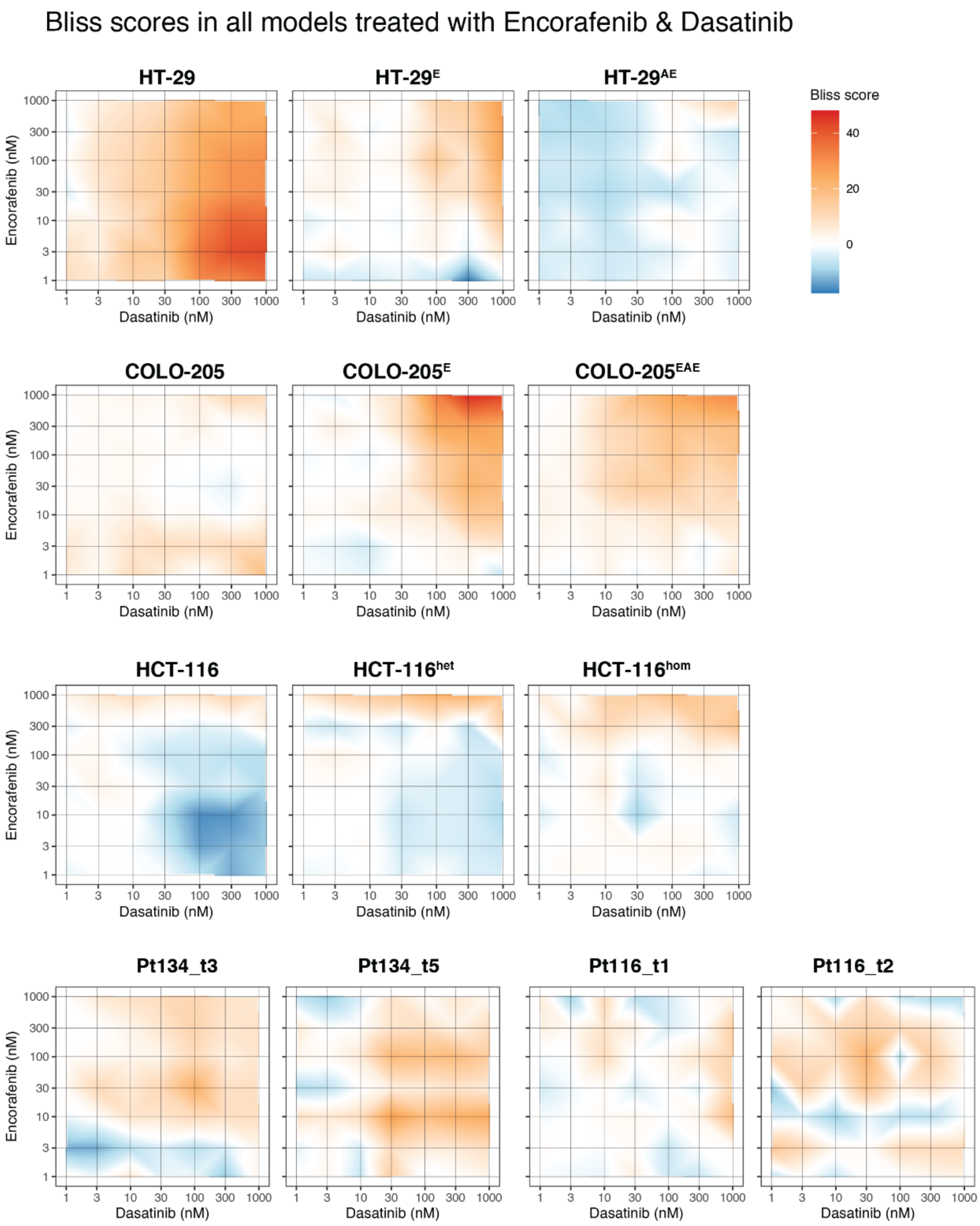


**Supplementary Figure 12. Bliss synergy landscapes for combined BRAF and SRC inhibitor dasatinib across cell line and PDOs.**

Dose-response heatmaps showing Bliss scores for the combination of encorafenib and dasatinib across parental and resistant HT-29 and COLO-205 sublines, isogenic HCT116 models, and selected PDOs. Bliss scores represent deviations from Bliss independence, with positive values indicating synergistic interactions and negative values indicating antagonism. Bliss score surfaces were interpolated (akima::interp, R) on log10 concentrations for visualization.

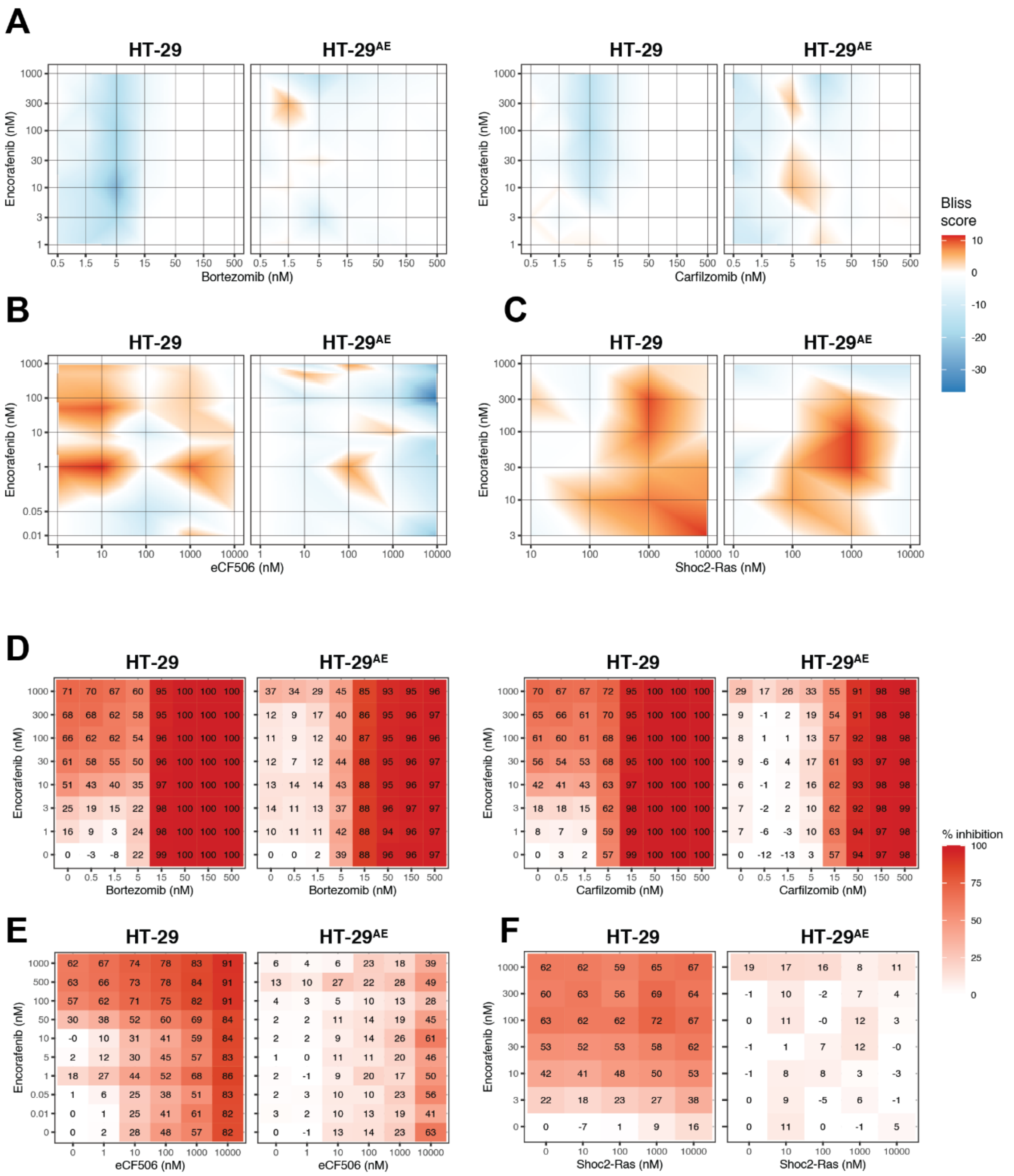


**Supplementary Figure 13. Impact on viability and synergy of encorafenib combined with proteasome, SRC and SHOC2-RAS inhibition in HT-29 parental and resistant sublines.** Dose-response heatmaps showing Bliss scores (**A-C**) and percentage viability inhibition (**D-F**) for the combination of encorafenib with proteasome inhibitors bortezomib and carfilzomib (**A** and **D**), SRC inhibitor eCF506 (**B** and **E**) and the SHOC2-RAS interaction inhibitor SHOC2–RAS PPI-IN-1 (**C** and **F**) in parental HT-29 cells and in the encorafenib/afatinib–resistant subline HT-29_10AE. Bliss scores represent deviations from Bliss independence, with positive values indicating synergy and negative values indicating antagonism. Bliss score surfaces were interpolated (akima::interp, R) on log10 concentrations for visualization.

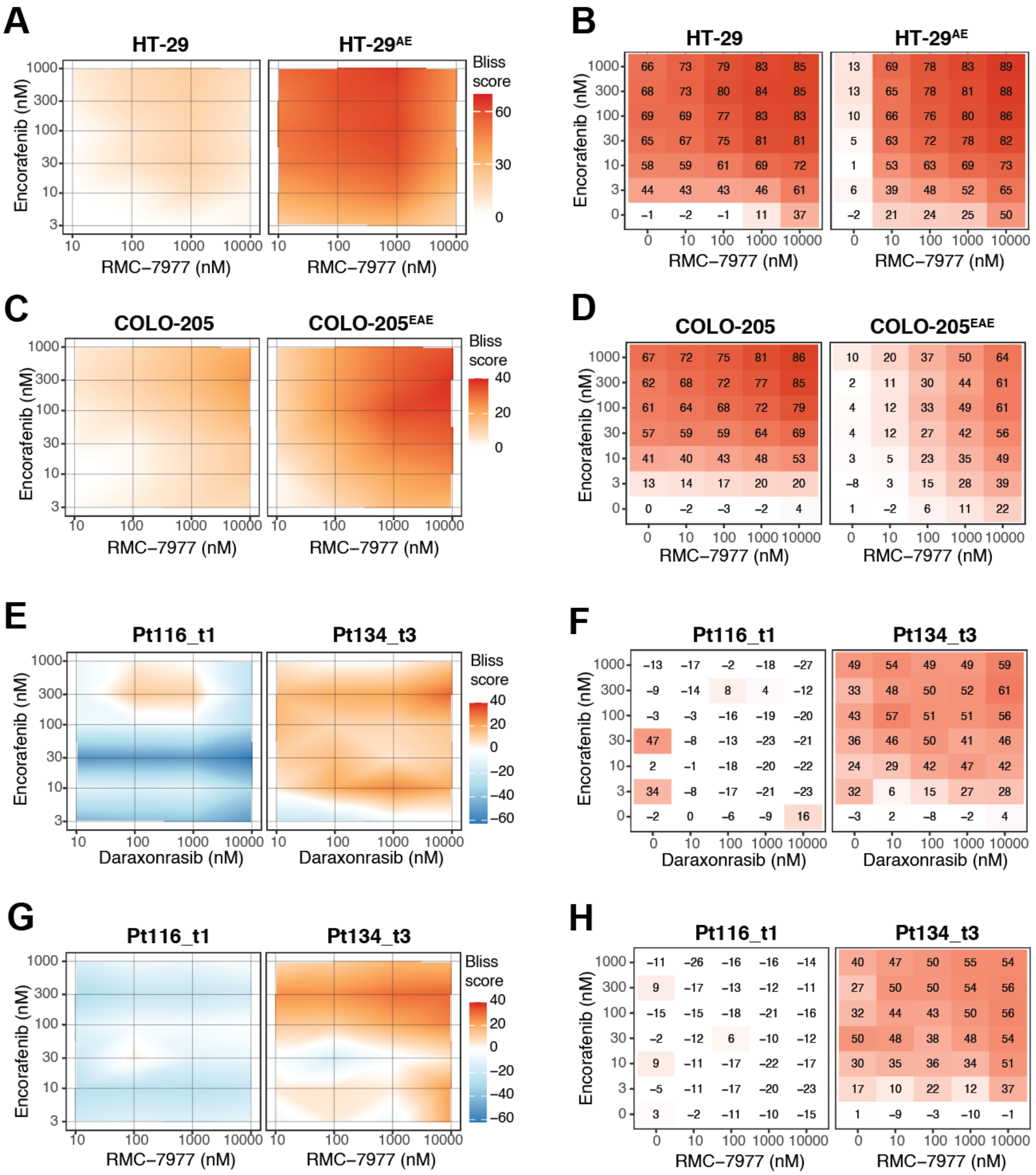


**Supplementary Figure 14. Impact on viability and synergy of encorafenib combined with RAS(ON) inhibitors cell lines and PDOs.**

Heatmaps showing Bliss scores (**A, C, E,** and **G**) and percentage viability inhibition (**B, D, F** and **H**) for the combination of encorafenib with RAS inhibitors daraxonrasib and RMC-7977 in cell lines (**A**-**D**) and PDOs (**E**-**H**).

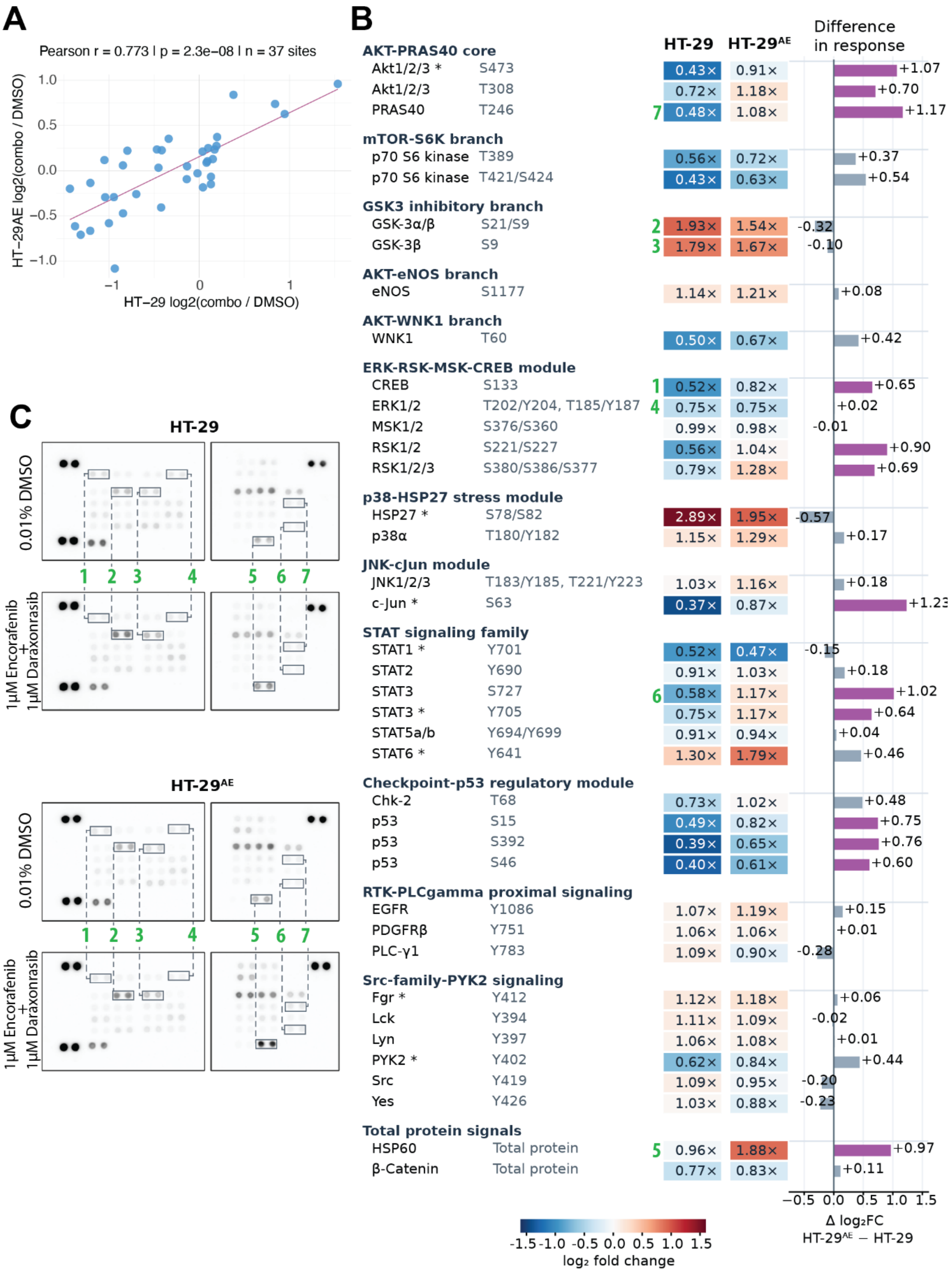
A
Pearson r = 0.773 | p = 2.3e−08 | n = 37 sites
HT−29AE log2(combo / DMSO)
HT−29 log2(combo / DMSO)
B
HT-29
HT-29AE
Difference in response
AKT-PRAS40 core
Akt1/2/3 * S473 0.43× 0.91× +1.07
Akt1/2/3 T308 0.72× 1.18× +0.70
PRAS40 T246 7 0.48× 1.08× +1.17
mTOR-S6K branch
p70 S6 kinase T389 0.56× 0.72× +0.37
p70 S6 kinase T421/S424 0.43× 0.63× +0.54
GSK3 inhibitory branch
GSK-3α/β S21/S9 2 1.93× 1.54× -0.32
GSK-3β S9 3 1.79× 1.67× -0.10
AKT-eNOS branch
eNOS S1177 1.14× 1.21× +0.08
AKT-WNK1 branch
WNK1 T60 0.50× 0.67× +0.42
ERK-RSK-MSK-CREB module
CREB S133 1 0.52× 0.82× +0.65
ERK1/2 T202/Y204, T185/Y187 4 0.75× 0.75× +0.02
MSK1/2 S376/S360 0.99× 0.98× -0.01
RSK1/2 S221/S227 0.56× 1.04× +0.90
RSK1/2/3 S380/S386/S377 0.79× 1.28× +0.69
p38-HSP27 stress module
HSP27 * S78/S82 2.89× 1.95× -0.57
p38α T180/Y182 1.15× 1.29× +0.17
JNK-cJun module
JNK1/2/3 T183/Y185, T221/Y223 1.03× 1.16× +0.18
c-Jun * S63 0.37× 0.87× +1.23
STAT signaling family
STAT1 * Y701 0.52× 0.47× -0.15
STAT2 Y690 0.91× 1.03× +0.18
STAT3 S727 6 0.58× 1.17× +1.02
STAT3 * Y705 0.75× 1.17× +0.64
STAT5a/b Y694/Y699 0.91× 0.94× +0.04
STAT6 * Y641 1.30× 1.79× +0.46
Checkpoint-p53 regulatory module
Chk-2 T68 0.73× 1.02× +0.48
p53 S15 0.49× 0.82× +0.75
p53 S392 0.39× 0.65× +0.76
p53 S46 0.40× 0.61× +0.60
RTK-PLCgamma proximal signaling
EGFR Y1086 1.07× 1.19× +0.15
PDGFRβ Y751 1.06× 1.06× +0.01
PLC-γ1 Y783 1.09× 0.90× -0.28
Src-family-PYK2 signaling
Fgr * Y412 1.12× 1.18× +0.06
Lck Y394 1.11× 1.09× -0.02
Lyn Y397 1.06× 1.08× +0.01
PYK2 * Y402 0.62× 0.84× +0.44
Src Y419 1.09× 0.95× -0.20
Yes Y426 1.03× 0.88× -0.23
Total protein signals
HSP60 Total protein 5 0.96× 1.88× +0.97
β-Catenin Total protein 0.77× 0.83× +0.11
-1.5 -1.0 -0.5 0.0 0.5 1.0 1.5
log2 fold change
Δ log2FC
HT-29AE − HT-29
C
HT-29
0.01% DMSO
1μM Encorafenib + 1μM Daraxonrasib
1 2 3 4 5 6 7
HT-29AE
0.01% DMSO
1μM Encorafenib + 1μM Daraxonrasib
1 2 3 4 5 6 7

**Supplementary Figure 15. Phosphorylation responses to combined BRAF and RAS inhibition in parental HT-29 and resistant HT-29$^{AE}$ cells.**

Cells were treated with encorafenib and daraxonrasib, each at 1 µM, or vehicle control (0.01% DMSO) for 72 h. (**A**) Correlation of treatment-induced phosphorylation changes between HT-29 and HT-29AE cells. Each point represents one of 37 phosphorylation-site measurements, expressed as $\log_2$ fold change relative to the corresponding DMSO control. The line indicates the fitted linear regression; Pearson's correlation coefficient and associated P value are shown. (**B**) Treatment responses grouped by signaling relationships and protein family. Heatmap colors represent $\log_2$ fold changes relative to the corresponding DMSO control, with blue indicating decreased signal and red indicating increased signal. Numerical labels indicate untransformed fold changes. HSP60 and β-catenin represent total-protein measurements and are displayed separately. Bars indicate differences in treatment response, calculated as $\Delta\log_2FC = \log_2FC(HT\text{-}29^{AE}) - \log_2FC(HT\text{-}29)$. Positive values indicate a higher treatment/control ratio in HT-29$^{AE}$ cells. Purple bars denote absolute differences $\geq\log_2(1.5)$, corresponding to a ≥1.5-fold difference between treatment/control ratios; remaining bars are gray. Asterisks identify targets with duplicate-spot coefficients of variation >25%. Green numbers identify the corresponding spots highlighted in panel C and Figure 5G. (**C**) Chemiluminescent images of the phospho-kinase arrays, with the two component membranes (parts A and B) shown on the left and right, respectively, for each condition. Each target is represented by duplicate capture-antibody spots. Boxes and dashed lines mark selected targets numbered in green: 1 - CREB; 2 - GSK-3α/β; 3 - GSK-3β; 4 - ERK1/2; 5 - HSP60; 6 - STAT3 (S727); and 7 - PRAS40.

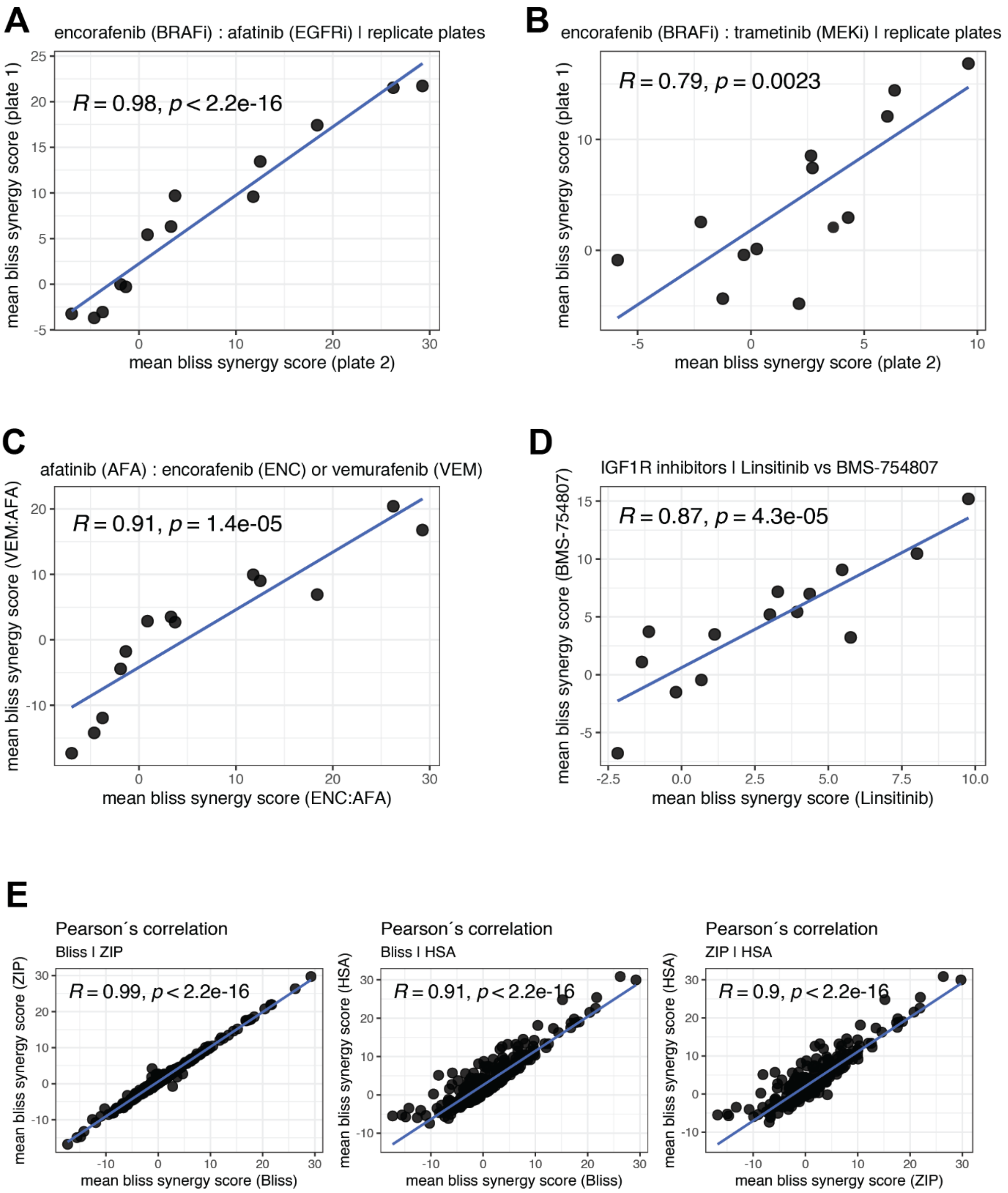


**Supplementary Figure 16. Robustness of synergy scoring across replicates and synergy models.**

Correlation of mean Bliss synergy scores between two independent biological replicates for the combination of encorafenib (BRAF inhibitor) and afatinib (EGFR inhibitor) **(A)** and for the combination of encorafenib and trametinib (MEK inhibitor) **(B)**. Correlation of mean Bliss synergy scores for afatinib combined with two BRAF inhibitors encorafenib or vemurafenib **(C)**, as well as encorafenib combined with two IGF1R inhibitors, linsitinib and BMS-754807 **(D)**, illustrating consistency across related drug classes. **(E)** Pairwise comparison of mean synergy scores calculated using Bliss independence (Bliss), Zero Interaction Potency (ZIP), and Highest Single Agent (HSA) models, showing Pearson correlation coefficients and corresponding p-values for each comparison. Each point represents an individual drug combination.
